# Can you run from your worries? The effects of exercise on anxiety-like behaviour and immune signaling in female and male mice

**DOI:** 10.64898/2026.04.08.717231

**Authors:** Madeleine Gene Maheu, Julia Mazur, Evelin Melekh, Madeleine King, Gurprince Attlas, Emma Cook, Sarah Bellaflor, Sunny F. Qureshi, Ahmad Mohammad, Shawn M Beaudette, Rebecca EK MacPherson, Paula Duarte-Guterman

## Abstract

Exercise is a positive health behaviour associated with improved mood. However, the mechanisms underlying the benefits of exercise on affective health are unclear, particularly with respect to type of exercise and sex. Chronic exercise decreases neuroinflammation, which is linked to improvements in mood and anxiety. However, exercise is also a physiological stressor that can transiently upregulate systemic inflammation, and its effects on neuroinflammation are not well understood. This study examined how acute and chronic exercise affect circulating and brain cytokine levels and anxiety-related behaviour in young healthy male and female mice. In Experiment 1, mice were placed on a treadmill for a two-hour bout of moderate exercise. Two hours after exercise, animals were either tested in the open field or euthanized for measurement of cytokines (IL-1β, TNF, IL-2, IL-4, IL-5, IL-6, IL-10, IL-12p70, IFN-γ, KC/GRO). In Experiment 2, mice underwent an 8-week moderate treadmill exercise paradigm followed by open field testing and tissue collection. Acute exercise decreased time spent in the centre of the open field in males only, suggesting increased anxiety-like behaviour in males. Acute exercise increased IL-6 and decreased TNF in serum, and increased amygdala principal component 1 (loading IL-12p70, IL-10, IFN-γ, and TNF) in both sexes. Chronic exercise increased open field centre entries, increased IL-6 in the prefrontal cortex, decreased TNF in the dorsal hippocampus, and had minimal effects on circulating cytokines in both sexes. These results demonstrate that the effects of exercise on anxiety-related behaviour and cytokine levels depend on recurrence, tissue, and brain region.

**NEW & NOTEWORTHY:** Our work highlights the contrast between anxiogenic and anxiolytic effects of acute versus chronic exercise, respectively, in healthy mice. Acute and chronic exercise differentially affected circulating and brain cytokines, providing insight into physiological adaptations to exercise. Both sexes demonstrated similar cytokine responses to exercise. These similarities are novel with respect to exercise research and noteworthy given sex differences in anxiety with respect to acute exercise.

## INTRODUCTION

The psychological benefits of exercise have been widely acknowledged and supported by anecdotal reports and extensive research. Regular exercise is associated with improved psychological well-being (1,2) and reduced anxiety in humans of either sex (3,4) and in male rodent models (5–8). However, the effects of acute exercise on anxiety-like behaviour in rodents have shown less consistent results (9–11) and have focused on male rodents. Acute versus chronic exercise may produce differential changes to brain and behaviour because of short versus long term physiological adaptations, which are not completely understood, especially in the brain (12). The limited research in this area is also hindered by lack of sex-comparisons, variability of exercise paradigms, and differences in behavioural testing methodology. These limitations make it challenging to form a concrete and encompassing conclusion pertaining to affective outcomes of short vs long term exercise regimen in animals.

The benefits of exercise on mood may arise, at least in part, from physiological processes involving the immune response to exercise. Cytokines are key signaling molecules in immune function that have also been implicated in the modulation of mood and anxiety (13). In humans, the capacity to synthesize and release cytokines into circulation is positively correlated with the severity of anxiety symptoms, suggesting that heightened inflammatory signaling may contribute to anxiety (13). In contrast, exercise induces a transient increase in cytokine release into circulation, which promotes an overall anti-inflammatory environment (14) and may help reduce anxiety over time (15). Generally, pro-inflammatory cytokines (such as IL-1β and TNF) can increase anxiety (16,17). Acute bouts of exercise can also transiently increase anxiety in humans (18), potentially due to short-term elevations in pro-inflammatory cytokines (14). However, with repeated exercise bouts, there is a gradual shift toward reduced systemic pro-inflammatory signaling and enhanced anti-inflammatory signaling (19–22). Together, this suggests that while acute exercise may transiently increase anxiety through pro-inflammatory activation, chronic exercise may have the opposite effect. Although the cytokine response to acute exercise has been well characterized peripherally (23–25), much less is known about the response in the brain. It is unclear whether the transient peripheral cytokine changes induced by acute exercise extends to the central nervous system. While chronic exercise can reduce neuroinflammation in disease models (26–30), it is unknown whether this anti-inflammatory effect remains relevant in healthy models and in both sexes. Previous work has demonstrated that inflammatory profiles and responses to stress and immune challenge can differ by brain region, with distinct differences between the prefrontal cortex, amygdala, and dorsal and ventral hippocampus (31–35). With respect to exercise, relatively little is known about brain regional cytokine responses. Existing studies in healthy mice have primarily focused on the hippocampus and indicate that acute exercise increases hippocampal IL-6 levels, decreases TNF levels, and has no effect on IL-1β levels (36,37). In contrast, chronic exercise increases hippocampal IL-1β in healthy rats (38). Findings seem to vary with exercise intensity, duration, experimental model, and potentially sex (36–38). Therefore, significant gaps remain in our understanding of regional cytokine responses to exercise, with few studies distinguishing between the dorsal and ventral hippocampus or examining the amygdala. In addition, the relationships between exercise-induced cytokine brain and serum changes and behavioural outcomes, such as anxiety, remain poorly understood, and sex differences have rarely been examined. Accordingly, the purpose of this study was threefold: (1) to determine the effects of acute and chronic exercise on anxiety-like behaviour; (2) to characterize the effects of acute and chronic exercise on various cytokines in the blood and functionally distinct brain regions; and (3) to determine whether there are any sex differences with respect to these measures in healthy adult mice.

## METHODS

### Animals and ethics

Female and male 8-wk-old C57BL/6J mice were purchased from The Jackson Laboratory (Bar Harbor, ME). Mice acclimatized for 7 days in the Brock University Comparative Biosciences Facility before commencing experimentation. All experimental groups were age-matched such that animals were the same age at the time of behavioural testing and tissue collection (Experiment 1 and 2 below). All mice were kept on a 12-h light:dark cycle and had ad libitum access to food and water through the entirety of the study. Experimental protocols were approved by the Brock University Animal Care Committee (AUP: #23-09-03) and compliant with the Canadian Council on Animal Care.

### Experimental design

#### Experiment 1: Acute Exercise

Mice were assigned to an acute exercise intervention with one subset of animals euthanized 2 h after the exercise bout (n=8/sex), and a second subset (n=10/sex) undergoing open field testing 2 h post exercise bout. These sample sizes were chosen based on previous work on the effects of exercise on circulating and brain cytokines conducted in male C57BL/6J mice (36,39–43). Baseline measures of all mice were taken immediately before group assignment to ensure no initial differences in body mass between experimental groups (sedentary males 31.83 ± 1.60 g; exercised males 31.83 ± 1.72 g; sedentary females 21.5 ± 1.31 g; exercised females 21.38 ± 0.74 g). A few days before running, all mice were acclimated to the treadmill for 10 minutes at 10 m/min and a 5% incline. Exercised mice ran for 120 min at a speed of 15 m/min and a 5% incline. Any mice that would not run were gently prodded on the rear with a nylon brush. Sedentary mice were placed on a non-moving treadmill replica in tandem with and for the same amount of time as the exercised group to control for the novel environment and for stress across groups.

#### Experiment 2: Chronic Exercise

Mice were assigned to either an exercise (n=8/sex) or sedentary (n=12/sex) group. We have previously reported significant behavioural and PNN content changes using a sample size of 12/group/sex (35). Four animals per sex in the exercise group were excluded prior to the start of the experiment due to a housing assignment error. Before assignment, body masses were taken, and all mice were habituated to the treadmill for two 10-minute sessions of low intensity running (10 m/min at a 5% incline) prior to assignment to experimental groups (sedentary males: 23.54 ± 1.59 g, exercised males: 23.34 ± 1.11 g, sedentary females: 17.38 ± 1.16 g, exercised females 17.91 ± 0.81 g). The chronic exercise paradigm utilized treadmill running for one hour per day, five days per week at an increasing speed and incline for eight weeks as done previously (35,44). This treadmill training protocol represents 65% VO_2_max and is moderate in intensity (39), with the intervention beginning at 20 m/min at 15 % incline and ending at 25 m/min at 25 % incline for the final two weeks. Sedentary controls were placed on a non-moving treadmill replica at the same time as the exercised group. During the final week of exercise training, open field testing was performed.

Mice that underwent the chronic exercise training were scanned using a dual energy X-ray absorptiometry (DXA; Scintica, Osteosys) to assess body composition, including lean mass, fat mass, bone mineral content, and bone mineral density, in anesthetized mice (vaporized isoflurane 5% in O_2_). Scans were performed one week prior to the start of the 8-week exercise intervention and immediately before euthanasia.

### Open field testing and analysis

Mice underwent open field testing 2 h after acute running (Experiment 1) and 2 hours post-exercise on the final two days of chronic running (Experiment 2). Mice were placed in an empty open-field testing arena with black walls and a white floor (40 cm x 40 cm x 40 cm) and allowed to explore for 5 min. Open field boxes were cleaned with Peroxigard solution between animals and all testing took place between 12 pm and 1 pm. All behavioural videos were recorded using an overhead GoPro (1920x1080 pixels; 30 fps), with animal tracking performed using DeepLabCut (DLC version 2.2.3.9). A custom DLC model was trained using user inputs designating the locations of four anatomical landmarks (nose, tail base, left and right ear) as well as arena boundaries (corners: top left, top right, bottom left, and bottom right). Specifically, 20 frames (*k*-means clustering) were extracted from 16 training videos, and the DLC model (ResNet50) was trained from these frames using the above-stated landmarks. Model training was performed over 200,000 epochs, resulting in a mean testing error of 3.74 pixels across all tracked positional landmarks. Once the model was trained and all behavioural videos were subsequently analyzed using the custom DLC model, with the tracked XY positional data derived from DLC being further analysed using custom MATLAB scripts (v2024a, The MathWorks Inc.). First, when animal tracking confidence was low (<0.9), the XY coordinate data were interpolated using a piecewise cubic Hermite interpolating polynomial. Next, positional landmarks were defined to distinguish between the arena periphery (within 10 cm of arena walls) and centre (beyond 10 cm of arena walls). To determine whether the mouse was in the periphery or centre of the arena, two anatomical landmarks (centre of head and tail base) had to cross the landmark boundary. Automated scoring included time spent in the centre, time spent in the corners, time spent immobile, average speed, as well as the number of entries into the centre of the arena. In addition to automated scoring, manual scoring methods were used to measure instances of rearing and grooming during open field testing by an experimenter blinded to the treatment conditions. Rearing was defined as any instance in which the mouse was standing on its hindlimbs. Rearing was further subdivided into the categories of “supported” or “unsupported”, with supported rearing being defined as hindlimb-standing against a wall and unsupported rearing defined as freely standing on the hindlimbs without the support of a wall. Grooming was defined as any instance in which the mouse appeared to be licking its paws and/or rubbing its paws against its head or body.

### Tissue collection and preparation

Mice were anesthetized with vaporized isoflurane (5%) to the extent of pedal reflex loss. Mice were euthanized via exsanguination through blood draw directly from the left ventricle. Blood samples were stored at 4°C for 24 h, centrifuged for 10 min at 1500 RCF at 4°C, and serum stored at –80°C. Adrenal glands were dissected and weighed as an indicator of stress. Brains were extracted, frozen on dry ice, and stored at –80°C for subsequent brain punches. Brains were mounted and embedded in OCT compound, and coronally sliced at a thickness of 300 μm using a cryostat. Distinct brain regions (prefrontal cortex, hippocampus, amygdala) were dissected, and the dorsal and ventral hippocampus were analyzed separately to account for their differential functions (45). Frozen sections were mounted onto microscope slides and the prefrontal cortex (Bregma 2.92 to 2.22), dorsal hippocampus (Bregma –1.22 to –2.80), ventral hippocampus (Bregma –2.92 to –3.88) and amygdala (Bregma -0.96 to -2.70) were quickly punched out of sections from both hemispheres. Punches for the regions of interest were stored at -80°C until they were homogenized in Invitrogen cell lysis buffer II, supplemented with phenylmethylsulphonyl fluoride (PMSF) and protease inhibitor (Thermo Fisher Scientific). Samples underwent homogenization in a FastPrep homogenizer at a speed of 6.5 for 45 s. The brain homogenates were placed on a rocker at 4°C for 20 min before being centrifuged at 10,000 g for 15 min at 4°C and supernatant was collected as stored at –80°C. Protein concentration was determined in triplicates using a bicinchoninic acid assay and homogenates were stored at –80°C.

### Multiplex immunoassay for cytokines

To determine the cytokine content in serum and brain tissue, a multiplex immunoassay was conducted using the V-PLEX Pro-Inflammatory Mouse Panel-1 Kit (IL-1β, TNF, IL-2, IL-4, IL-5, IL-6, IL-10, IL-12p70, IFN-γ, KC/GRO; Meso Scale Discovery, Rockville, MD). Samples of prefrontal cortex, and dorsal and ventral hippocampus were run in duplicate. Amygdala samples were run in singlet. 50 µL of diluted sample (1:2 for serum and 1:1.5 for brain tissue) was added to each well of a 96 well plate, which was then sealed and incubated for 24 hours at 4°C. Wells were washed with wash buffer and 25 µL of detection antibody solution was added to each well. Wells were sealed and left to incubate for 2 h at room temperature. Wells were washed again and 150 µL of read buffer was added to each well. Plates were read using a Meso Scale Discovery Quickplex SQ 120 MM instrument. Analyses were done using a calibration curve of known standards to which data were fitted to determine cytokine concentrations using the MSD software (Discovery Workbench). Cytokine levels were made relative to the total protein content loaded per well.

### Statistical analyses

GraphPad Prism (version 8.4.3.686) was used to run all analyses of variance (ANOVAs). Morphometric data were subject to log or box-cox transformation when assumptions of normality and/or homogeneity of the variance were violated. Observations more than three standard deviations from the mean were removed as outliers. ANOVAs using sex (female, male) and treatment (sedentary, exercised) as between subject factors were performed on post-morphometric measures, adrenal to body mass ratio, and behavioural data. Given previous work in humans on the effects of exercise on IL1β, IL-6, and TNF (14,23–25), ANCOVAs were performed on these individual cytokines using plate as a covariate, as samples were distributed over two plates. In the event of a significant interaction, post-hoc tests were used and subjected to Tukey correction. Significance was identified as p ≤ 0.05 and trends identified when 0.05 < p ≤ 0.08.

To provide an exploratory analysis of various cytokines, data from the ten cytokines were reduced and extracted using principal component analysis (PCA) using SPSS version 30. All cytokine data underwent box-cox transformation before principal component analyses were conducted. A Kaiser-Meyer-Olkin test for sampling adequacy was performed for each PCA, and this threshold value was set to 0.6 to continue the analysis. Extracted components were subjected to varimax rotation. Separate PCAs were run for each experiment and tissue (prefrontal cortex, dorsal hippocampus, ventral hippocampus, amygdala, and serum) for a total of ten PCAs. For an objective approach to data reduction, only factors explaining ≥10% of the cytokine variation as measured by the eigenvalues (≥ 1) were reported and included in subsequent analyses. Factors that met these criteria were further analyzed using ANOVAs. For cases in which cytokines were not detectable, values were inputted with half of the lowest level of detection. However, IL-12 and IL-4 were excluded from serum analyses because over half of samples contained undetectable amounts of these cytokines. Lack of detectability was not related to exercise treatment or sex.

## RESULTS

### Experiment 1: Behavioural and biological changes with acute exercise

#### Acute exercise decreased average speed and increased time spent immobile

We determined whether acute exercise or sex influenced anxiety-like behaviour by examining differences across multiple measures in the open field, including supported and unsupported rearing, time grooming, average speed, time immobile, time in arena centre, centre entries, and time in corners (Figure 1). Exercise affected the percent time spent in the centre of the open field (Figure 1A) differently in females and males (interaction: F (1, 36) = 5.407, p = 0.0258), with sedentary males spending more time in the centre than exercised males (*p* = 0.0572) and sedentary females (*p* = 0.0133). Time spent in the corners of the arena (Figure 1B) was not affected by exercise (main effect of exercise: F (1, 36) = 1.663, p = 0.2054, sex by exercise interaction: F (1, 36) = 1.360, p = 0.2512) but females spent more time in the corners than males (effect of sex: F (1, 36) = 9.667, p = 0.0037). Entries into the centre of the arena (Figure 1C) did not change with exercise (F (1, 36) = 0.6686, p = 0.4189), sex (F (1, 36) = 0.08747, p = 0.7691), or their interaction (F (1, 36) = 1.748, p = 0.2542).

**Figure 1.**
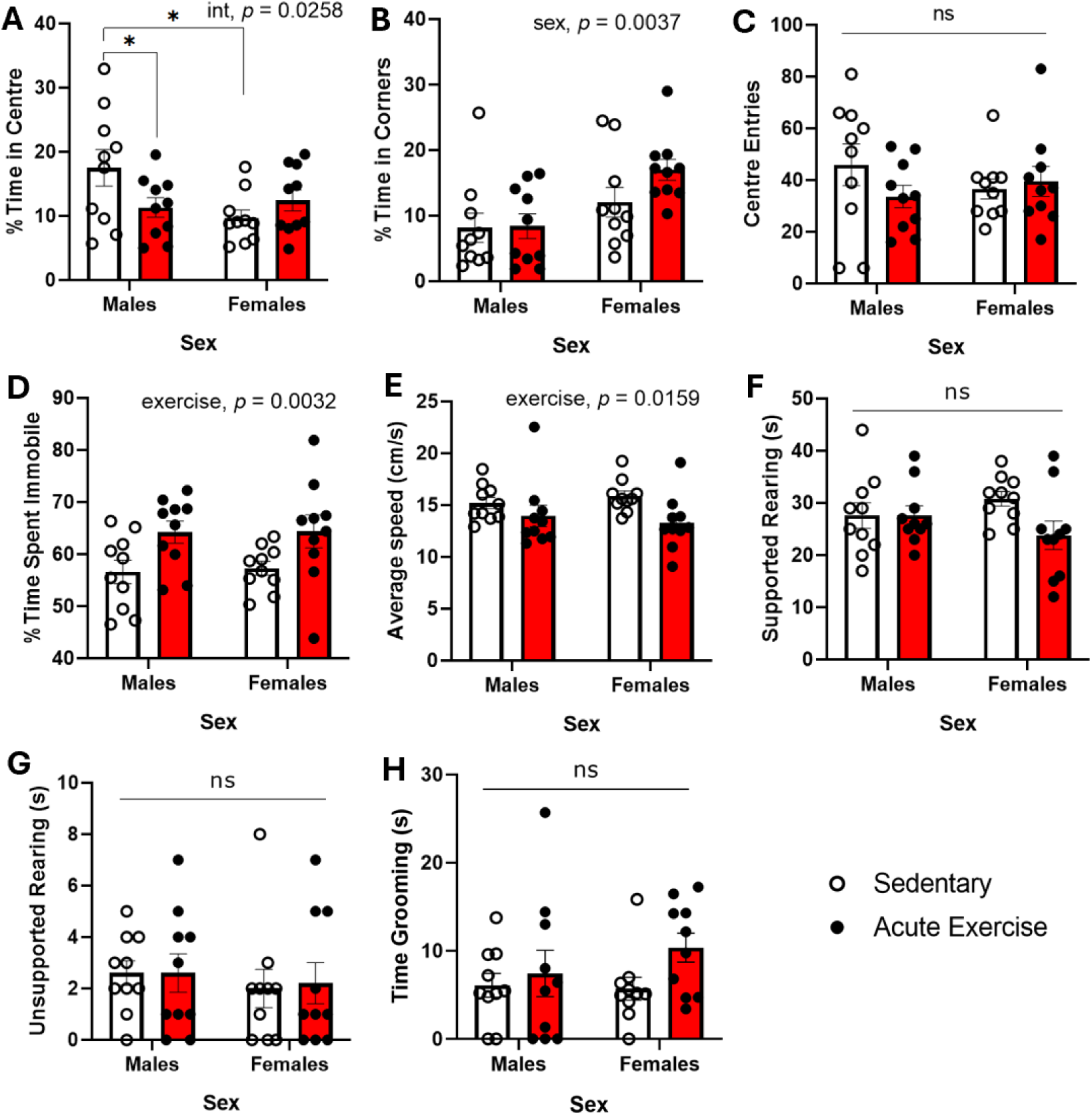
Effects of acute exercise and sex on locomotion and location during the open field test. Sedentary males spent (A) more time in the centre and (B) females spent more time in the corners of the arena than males (p < 0.01). (C) There was no effect of exercise, sex, or interaction on centre entries (p’s > 0.05). Exercised animals (D) spent more time immobile and (E) moved at a slower average speed than sedentary controls (*p’s* < 0.05). There were no exercise, sex or interaction effects for (F) supported rearing, (G) unsupported rearing, and (H) time spent grooming. Error bars indicate standard error. Note, \**p* < 0.05. n = 10 per group and sex.

For percent time spent immobile (Figure 1D), there was an effect of exercise, favouring immobility among the exercised animals (F (1, 36) = 10.01, p = 0.0032), no effect of sex (F (1, 36) = 0.03626, p = 0.8501), and no interaction (F (1, 36) = 0.01315, p = 0.9094). Similarly, for average speed, there was a significant effect of exercise (F (1, 36) = 6.405, p = 0.0159), with exercised animals moving slower than sedentary controls (Figure 1E). There was no effect of sex (F (1, 36) = 2.295e-006, p = 0.9988), and no interaction (F (1, 36) = 0.7984, p = 0.3375).

There was no effect of acute exercise or sex on supported rearing (exercise: F (1, 36) = 2.609, p = 0.1150), sex: (F (1, 36) = 0.01917, p = 0.8906), interaction: (F (1, 36) = 2.609, p = 0.1151; Figure 1F)), unsupported rearing (exercise: (F (1, 36) = 0.02018, p = 0.8878), sex: (F (1, 36) = 0.5045, p = 0.4821), interaction: (F (1, 36) = 0.02018, p = 0.8878; Figure 1G)), or time spent grooming (exercise: (F (1, 36) = 2.236, p = 0.1435), sex: (F (1, 36) 63 = 0.8132, p = 0.3732), interaction: (F (1, 36) = 1.09, p = 0.3028; Figure 1H)).

#### Changes in serum IL-1 β, IL-6, and TNF after acute exercise

We examined the effects of acute exercise on certain individual cytokines, IL-1β, IL-6, and TNF in serum (Figure 2A-C). We found that exercise increased circulating levels of IL-6 in both sexes (effect of exercise: F (1, 27) = 4.681, p = 0.0395; exercise by sex interaction was not significant: F (1, 27) = 0.2516, p = 0.6200) and males having significantly greater serum IL-6 levels than females (main effect of sex: F (1, 27) = 6.066, p = 0.0204). Acute exercise decreased circulating levels of TNF (F (1, 28) = 4.212, p = 0.050; exercise by sex interaction was not significant (F (1, 28) = 0.702, p = 0.410), with a trend for females having lower TNF than males (F (1, 28) = 3.399, p = 0.076). For IL-1β, there was no effect of exercise (F (1, 28) = 0.171, p = 0.683)), sex (F (1, 28) = 0.232, p = 0.634), or interaction (F (1, 28) = 0.258, p = 0.615) on serum levels of IL-1β.

**Figure 2.**
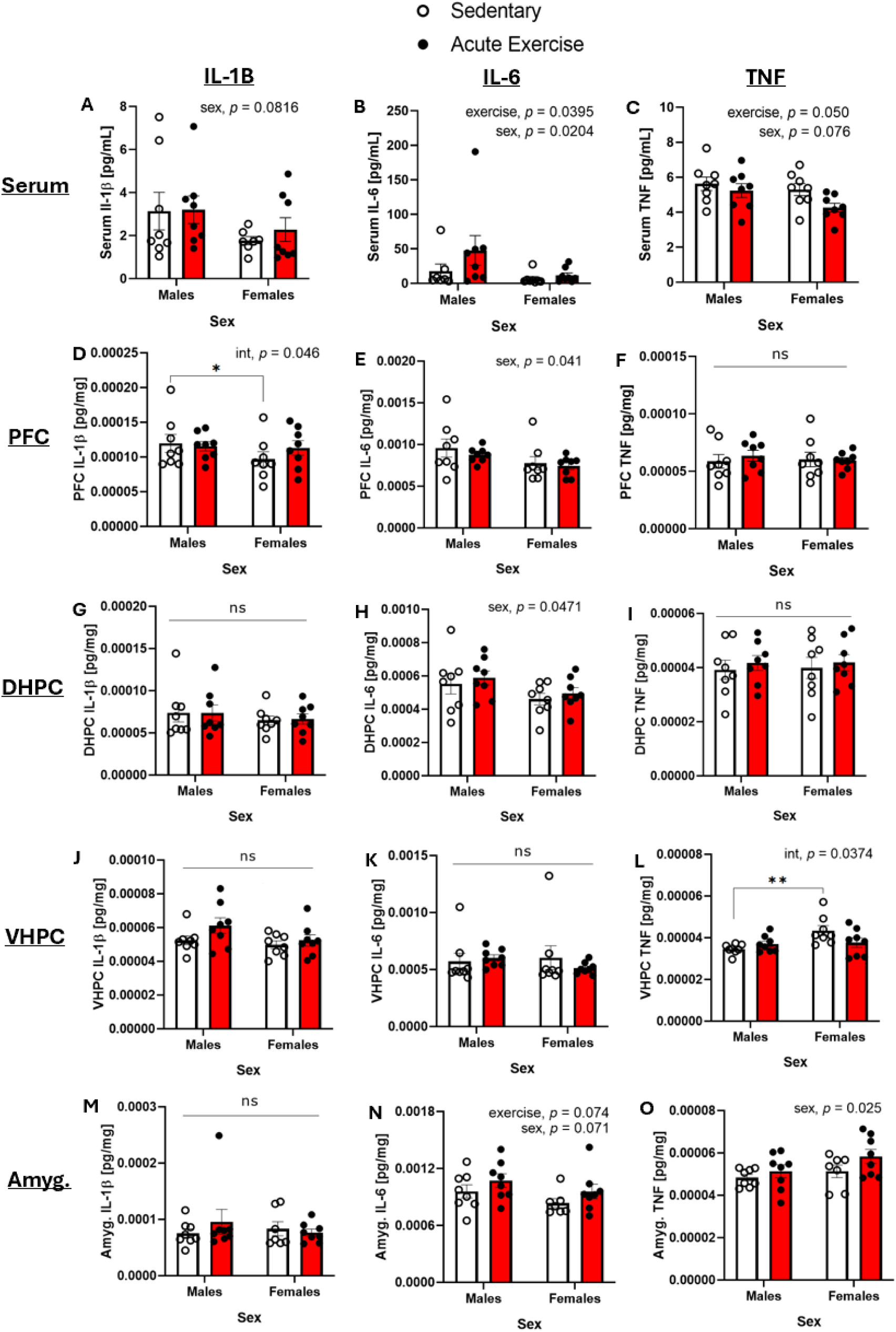
Effect of acute exercise and sex on IL-1β, IL-6 and TNF. Serum levels of (A) IL-1β, (B) IL-6, and (C) TNF. Prefrontal cortical levels of (D) IL-1β, (E) IL-6) and (F) TNF. Dorsal hippocampal levels of (G) IL-1β, (H) IL-6, and (I) TNF. Ventral hippocampal levels of (J) IL-1β, (K) IL-6, and (L) TNF. Amygdala levels of (M) IL-1β, (N) IL-6, and (O) TNF. Concentration levels were adjusted to protein concentration and are expressed as pg. cytokine/mg protein. Error bars indicate standard error. Note, * *p* < 0.05, ** *p* < 0.01. PFC: prefrontal cortex, DHPC: dorsal hippocampus, VHPC: ventral hippocampus, Amyg: Amygdala. n = 7-8 per group and sex.

#### Specific changes in brain cytokines after acute exercise

In the prefrontal cortex (Figure 2D-F), There was an interaction between acute exercise and sex on IL-1β content (*F* (1, 27) = 4.379, *p* = 0.046). Specifically, sedentary females had significantly less IL-1β in the prefrontal cortex than sedentary males (*p* = 0.011), but this difference was not found between the exercised females and exercised males (*p* = 0.801). Males had greater amounts of IL-6 in the prefrontal cortex than females (main effect of sex: *F* (1, 28) = 6.730, *p* = 0.041), but there was no effect of acute exercise on prefrontal IL-6 (main effect of exercise: *F* (1, 28) = 1.850, *p* = 0.185; interaction: *F* (1, 28) = 0.1134, *p* = 0.7388). TNF levels were not affected by acute exercise or sex in the prefrontal cortex (exercise: *F* (1, 28) = 0.1415, *p* = 0.7096, sex: F (1, 28) = 0.06702, *p* = 0.7976), interaction: *F* (1, 28) = 0.3354, *p* = 0.5671).

In the dorsal hippocampus (Figure 2G-I), acute exercise did not affect levels of IL-1β (exercise: *F* (1, 28) = 0.007602, *p* = 0.9311, sex: *F* (1, 28) = 0.9379, *p* = 0.3411, interaction: *F* (1, 28) = 0.01429, *p* = 0.9057), IL-6 (exercise: *F* (1, 28) = 0.6323, *p* = 0.4332, interaction: *F* (1, 28) = 3.447e-005, *p* = 0.9954) or TNF (exercise: *F* (1, 28) = 0.4662, *p* = 0.5004, sex: *F* (1, 28) = 0.01519, *p* = 0.9028, interaction: *F* (1, 28) = 0.007668, *p* = 0.9308). However, we found sex differences in IL-6 levels in the dorsal hippocampus, with males demonstrating higher concentrations than females (main effect of sex: F (1, 28) = 4.312, *p* = 0.0471; Figure 2H).

In the ventral hippocampus (Figure 2J-L), there was an interaction between acute exercise and sex for TNF levels (*F* (1, 28) = 4.776, *p* = 0.0374) with sedentary males demonstrating lower TNF levels than sedentary females (*p* = 0.0047), but this difference was not found between the exercised males and females (*p* = 0.9604; Figure 2L). Acute exercise did not affect ventral hippocampal levels of IL-1β (exercise: *F* (1, 28) = 2.863, *p* = 0.1017, sex: (1, 28) = 2.951, *p* = 0.096, interaction: *F* (1, 28) = 0.8402, *p* = 0.3672) or IL-6 (exercise: *F* (1, 28) = 0.2125, *p* = 0.6484, sex: *F* (1, 28) = 0.1833, *p* = 0.6719, interaction: *F* (1, 28) = 0.8720, *p* = 0.3584).

In the amygdala (Figure 2M-O), acute exercise did not affect IL-1β levels (*F* (1, 26) = 0.2092, *p* = 0.6511), sex (*F* (1, 26) = 3.273, *p* = 0.0816) or interaction (*F* (1, 26) = 0.1220, *p* = 0.7296). For IL-6, there was a trend for a main effect of acute exercise (*F* (1, 26) = 3.464, p = 0.074), and sex (*F* (1, 26) = 3.553, *p* = 0.071; Figure 2N), but no significant interaction (*F* (1, 26) = 0.126, *p* = 0.726). For TNF, females had higher concentrations in the amygdala compared to males (main effect of sex: *F* (1, 26) = 5.642, *p* = 0.025), but no effect of acute exercise (*F* (1, 26) = 2.881, *p* = 0.102), nor an interaction with sex (*F* (1, 26) = 0.00, *p* = 1.00; (Figure 2O).

#### Characterization of patterns in serum cytokines after acute exercise

A principal component analysis (Supplementary Figure 1A) of all cytokines, except IL-4 and IL-12p70 (as these cytokines were not consistently detected in serum), revealed that three factors accounted for 66.10% of cytokine variation observed from serum samples (Acute Serum Factor 1, 36.77%, Acute Serum Factor 2, 16.21%; Acute Serum Factor 3, 13.12%). To test for exercise and sex-dependent differences in the mean values of these extracted components, three separate ANOVAs were performed on Acute Serum Factor 1, 2, and 3. For Acute Serum Factor 1 (Supplementary Figure 1B), there was no effect of exercise (*F* (1, 28) = 0.4939, *p* = 0.4880) or interaction (*F* (1, 28) = 0.04764, *p* = 0.8288), but there was a significant effect of sex, with females demonstrating lower values of Acute Serum Factor 1 compared to males (*F* (1, 28) = 6.964, *p* = 0.0134; driven by high loadings of KC/GRO, IL-6, and moderate IL-10 and IL-1β). For Acute Serum Factor 2 (Supplementary Figure 1C), there was an effect of exercise, with acutely exercised animals demonstrating lower values of Acute Serum Factor 2 compared to sedentary animals (*F* (1, 28) = 4.632, *p* = 0.0401; driven by high loadings of IL-2 and moderate IFN-γ, and IL-5), no other differences were detected (no effect of sex (*F* (1, 28) = 8.738e-005, *p* = 0.9926) or interaction (*F* (1, 28) = 1.169, *p* = 0.2887). For Acute Serum Factor 3 (Supplementary Figure 1D), there was a significant interaction between acute exercise and sex (*F* (1, 28) = 4.337, *p* = 0.0465), however, post-hoc tests did not detect significant differences between any of the groups (all *p’s >* 0.1995).

#### Characterization of patterns in brain cytokines after acute exercise

In the prefrontal cortex, a principal component analysis of all cytokine concentrations revealed that two factors accounted for 76.75% of biomarker variation (Acute PFC Factor 1, 64.63%; Acute PFC Factor 2, 12.12%). All loadings for both extracted factors were positive loadings (Supplementary Figure 2A). For Acute PFC Factor 1 (Supplementary Figure 2B) and Acute PFC Factor 2 (Supplementary Figure 2C), there were no effects of exercise (F (1, 27) = 0.03708, p = 0.8487; F (1, 27) = 1.740, p = 0.1982, respectively), sex (F (1, 27) = 1.741, p = 0.1981; F (1, 27) = 0.2009, p = 0.6576, respectively), or interaction (F (1, 27) = 0.005447, p = 0.9417; F (1, 27) = 0.5292, p = 0.4732, respectively).

Analysis of dorsal hippocampal cytokine concentrations using principal component analysis (Supplementary Figure 3A-D) revealed three principal factors that together explained 69.78% of the overall variance (Acute DHPC Factor 1: 44.12%; Acute DHPC Factor 2: 13.89%; Acute DHPC Factor 3: 11.77%). There was no effect of exercise, sex, or interaction for Acute DHPC Factor 1 (exercise: (F (1, 28) = 0.6780, p = 0.4172), sex: (F (1, 28) = 0.3805, p = 0.5423), interaction: (F (1, 28) = 0.3824, p = 0.5413) or Acute DHPC Factor 3 (exercise: (F (1, 28) = 2.185, p = 0.1505), sex: (F (1, 28) = 1.036, p = 0.3175), interaction: (F (1, 28) = 1.226, p = 0.2776). For Acute DHPC Factor 2, there was also no effect of exercise (F (1, 28) = 0.1654, p = 0.6873) or interaction (F (1, 28) = 0.1625, p = 0.6899), but there was a trend for a main effect of sex (Supplementary Figure 3C), with females showing lower measures of Acute DHPC Factor 2 than males (F (1, 28) = 3.528, p = 0.0708).

In the ventral hippocampus, principal component analysis of cytokine concentrations (Supplementary Figure 4A-E) revealed that four factors accounted for 74.92% of biomarker variation (Acute VHPC Factor 1, 35.03%, Acute VHPC Factor 2, 16.53%, Acute VHPC Factor 3, 13.19%, Acute VHPC Factor 4, 11.17%). There was no effect of exercise, sex, or interaction for Acute VHPC Factor 1 (exercise: (*F* (1, 28) = 0.4487, p = 0.5084), sex: (*F* (1, 28) = 0.05975, *p* = 0.8087), interaction: (*F* (1, 28) = 0.9166, *p* = 0.3466)), Acute VHPC Factor 2 (exercise: (F (1, 28) = 0.06033, *p* = 0.8078), sex: (*F* (1, 28) = 2.302, *p* = 0.1404), interaction: (*F* (1, 28) = 1.721, p = 0.2002)), Acute VHPC Factor 3 (exercise: (*F* (1, 28) = 1.109, *p* = 0.3012), sex: (*F* (1, 28) = 3.139, *p* = 0.0873), interaction: (F (1, 28) = 2.940, *p* = 0.0974)), or Acute VHPC Factor 4 (exercise: (*F* (1, 28) = 0.0009444, *p* = 0.9757), sex: (*F* (1, 28) = 1.810, *p* = 0.1893), interaction: (F (1, 28) = 1.716, *p* = 0.200).

Principal component analysis of cytokine concentrations within the amygdala (Supplementary Figure 5A-D) indicated that three components together explained 73.23% of the total variance (Acute Amygdala Factor 1, 39.99%, Acute Amygdala Factor 2, 22.23%, Acute Amygdala Factor 3, 11.02%). For Acute Amygdala Factor 1, there was an effect of exercise (*F* (1, 27) = 5.829, *p* = 0.0228), with exercised animals demonstrating higher measures of Acute Amygdala Factor 1 (driven by high loadings of IL-12p70, IL-10, IFN-γ, and TNF; Supplementary Figure 5B), irrespective of sex (main effect of sex: *F* (1, 27) = 0.001835, *p* = 0.9661). Although the exercise effect appeared to be driven by the exercised females, the interaction effect was not significant (*F* (1, 27) = 2.941, *p* = 0.0978). For Acute Amygdala Factor 2 and Acute Amygdala Factor 3, there was no effect of exercise (F (1, 27) = 0.0001036, *p* = 0.9920; *F* (1, 27) = 0.9340, *p* = 0.3424, respectively), sex (F (1, 27) = 0.2799, *p* = 0.6011; *F* (1, 27) = 0.0007299, *p* = 0.9786, respectively), or interaction (*F* (1, 27) = 2.279, *p* = 0.1427; *F* (1, 27) = 0.002924, *p* = 0.9573, respectively).

### Experiment 2: Biological and behavioural changes after chronic exercise

#### Endurance trained females demonstrated greater changes in bone mineral density

All changes in morphometric parameters with exercise and sex are summarized in Table 1. Exercised animals showed greater increases in lean mass over the eight-week intervention compared to sedentary animals (F (1, 36) = 4.207, *p* = 0.0476; Table 1), and although this effect seemed to be driven by the exercised females, the interaction was not significant (F (1, 36) = 2.518, *p* = 0.1213). Exercised females had significantly greater increases in bone mineral density compared to sedentary females (*p* = 0.0469; interaction between sex and exercise (F (1, 36) = 5.269, *p* = 0.0276). There was a significant interaction between sex and exercise for lean and fat percentage body composition (F (1, 36) = 7.354, *p* = 0.0102), but follow-up multiple comparison tests showed no meaningful differences between sedentary and exercised females, nor sedentary and exercised males. Regardless of exercise, we found significant sex differences in body and bone composition across nearly all morphometric parameters (Table 1).

**Table 1.** Effects of chronic exercise training and sex on morphometric parameters and adrenal mass in male and female mice.

|  | Male |  | Female |  | ANOVA, *p* |  |  |
| --- | --- | --- | --- | --- | --- | --- | --- |
|  | Sed. | Ex. | Sed. | Ex. | Ex. | Sex | Intx |
| Body mass (g) | 27.45±2.67 | 28.475±1.87 | 21.05±0.905 | 20.91±0.93 | 0.5038 | **<0.0001** | 0.8172 |
| Change in body mass | 3.91±4.25 | 5.14±2.98 | 3.68±2.06 | 3±1.74 | 0.7781 | 0.2324 | 0.3357 |
| Fat mass (g) | 4.36±0.58 | 4.361±0.36 | 2.65±0.21 | 2.87±0.22 | 0.1728 | **<0.0001** | 0.2353 |
| % Fat mass | 15.37±0.94 | 14.807±0.65 | 12.57±0.83 | 13.44±0.74 | 0.5675 | **<0.0001** | **0.0102** |
| Change in fat mass | 1.42±1.17 | 1.39±0.82 | 0.56±0.51 | 0.99±0.40 | 0.4540 | **0.0229** | 0.3889 |
| Change in fat % | 0.12±1.07 | 0.25±1.235 | -0.73±1.492 | -1.56±1.163 | 0.6039 | **0.0567** | 0.4824 |
| Lean mass (g) | 23.92±2.03 | 25.07±1.461 | 18.44±0.99 | 18.486±0.58 | >0.1978 | **<0.0001** | 0.2639 |
| % lean mass | 84.63±0.94 | 85.19±0.65 | 87.43±0.83 | 86.56±0.737 | 0.5675 | **<0.0001** | **0.0102** |
| Change in lean mass (g) | 7.58±4.02 | 7.621±3.85 | 4.59±2.73 | 7.86±0.83 | **0.0476** | 0.4370 | 0.1213 |
| Change in lean % | 0.12±1.074 | 0.25±1.235 | -0.73±1.492 | -1.56±1.163 | 0.4540 | **0.0229** | 0.3889 |
| Adrenal/body mass % | 0.014±0.007 | 0.012±0.005 | 0.020±0.006 | 0.024±0.003 | >0.5066 | **<0.0001** | 0.1231 |
| Bone mineral density | 0.084±0.004 | 0.085±0.004 | 0.087±0.002 | 0.087±0.002 | 0.3588 | **0.0084** | 0.8951 |
| Change in bone mineral density | 0.003±0.006 | 0.001±0.006 | -0.001±0.009 | 0.007±0.004 | 0.1300 | 0.5215 | **0.0277** |
| Bone mineral content | 0.43±0.046 | 0.44±0.035 | 0.35±0.015 | 0.38±0.013 | **0.0148** | **<0.0001** | 0.2907 |
| Change in bone mineral content | 0.036±0.085 | 0.078±0.027 | 0.048±0.029 | 0.053±0.018 | 0.1719 | 0.7175 | 0.2769 |
| Food intake (g) | 4.51±0.76 | 4.70±0.18 | 4.61±1.34 | 4.67±0.79 | 0.9409 | 0.8265 | 0.5682 |

Note: Data are presented as mean ± SD. Morphometric data were acquired via DXA scan at the start and end of the eight-week exercise regimen. Two-way ANOVAs were used to assess exercise and sex main effects and interaction.

#### Chronically exercised animals demonstrated greater entries into the centre of the open field

Chronically exercised animals demonstrated significantly greater number of entries into the centre of the open field (main effect of exercise: *F* (1, 36) = 10.46, *p* = 0.0026; Figure 3C)). However, percent time spent in the centre of the open field (Figure 3A) was not affected by exercise or sex (exercise: *F* (1, 36) = 1.249, *p* = 0.2711, sex: *F* (1, 36) = 0.001018, *p* = 0.9747), or interaction (*F* (1, 36) = 0.0004952, *p* = 0.9824) For percent time spent immobile during the open field test (Figure 3D), a significant interaction was found among the chronic exercise cohort, (*F* (1, 36) = 8.353, *p* = 0.0082), with sedentary females spending a significantly less time immobile than exercised females (*p* = 0.0134). For average speed during the open field test (Figure 3E), there was a significant interaction between chronic exercise and sex (*F* (1, 36) = 6.705, *p* = 0.0138), but follow-up analyses showed no meaningful significant differences between the groups. There were no significant chronic exercise, sex, or interaction effects on supported rearing (exercise, *F* (1, 36) = 0.3738, *p* = 0.5448; sex: (F (1, 36) = 0.2021, *p* = 0.6557; interaction: (F (1, 36) = 0.1167, *p* = 0.7346; Figure 3F) or unsupported rearing (exercise, (F (1, 36) = 0.5266, *p* = 0.4727, sex: F (1, 36) = 1.063, p = 0.3093, interaction: F (1, 36) = 2.954, *p* = 0.0943; Figure 3G). However, there was an effect of both chronic exercise and sex on grooming behaviour (Figure 3H), with exercised mice grooming less than sedentary controls (*F* (1, 36) = 4.487, *p* = 0.0411), and females grooming less than males (*F* (1, 36) = 6.207, *p* = 0.0175). Although chronically exercised females appear to be driving the exercise effect on grooming behaviour, the interaction was not significant (*p* = 0.5131).

**Figure 3.**
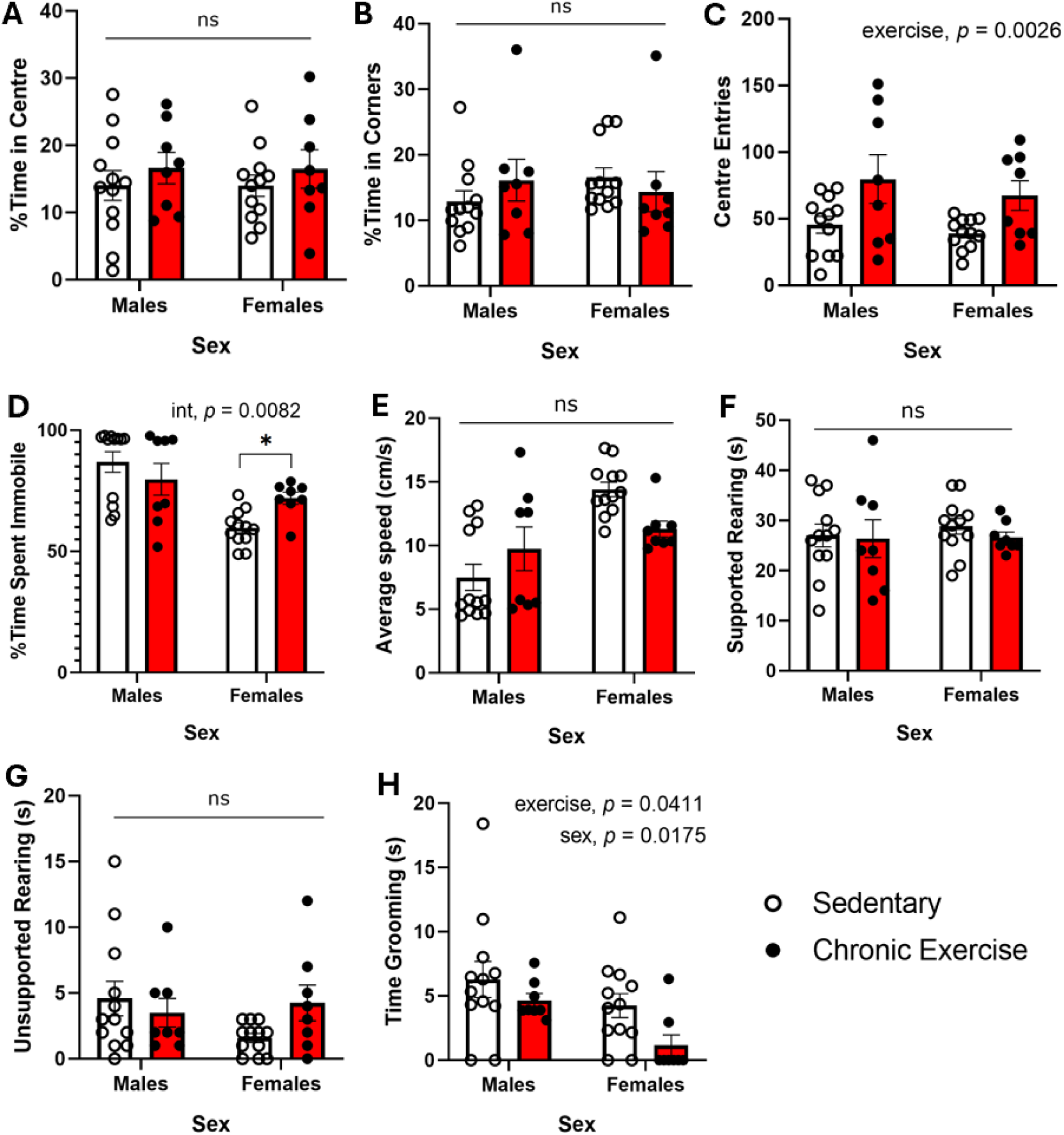
Locomotion and location among the chronic exercise cohort during the open field test. There were no exercise, sex or interaction effects for time spent in the (A) centre, and (B) corners (p’s > 0.05). (C) Exercised mice entered the centre more often than sedentary controls (*p* < 0.01). (D) Sedentary females spent less time immobile than exercised females (*p* < 0.05). (E) There was a significant interaction for average speed, but follow-up analyses showed no meaningful differences. There were no sex, exercise, or interaction effects for (F) supported or (G) unsupported rearing (p’s > 0.05). (H) Endurance-trained mice groomed less than sedentary controls, as did females (*p’s* < 0.05). Error bars indicate standard error. Note, \**p* < 0.05, \*\**p* < 0.01, \*\*\**p* < 0.001. n = 12 per sex in sedentary group and n = 8 per sex in exercised group.

#### Specific changes in serum cytokines after chronic exercise

In the serum (Figure 4A-C), exercise did not affect IL-1β concentration (exercise *F* (1, 36) = 0.01770, *p* = 0.8949), or interaction (*F* (1, 36) = 0.2365, p = 0.6297), but there was a main effect of sex (*F* (1, 36) = 4.084, *p* = 0.0508) with males having higher serum IL-1β than females (Figure 4A). Likewise, there was also an effect of sex on IL-6, with males having higher IL-6 levels than females (*F* (1, 34) = 27.19, *p* < 0.0001), but no effect of chronic exercise (*F* (1, 34) = 1.889, *p* = 0.548), nor an interaction (*F* (1, 34) = 0.1328, *p* = 0.254). For TNF, there was a significant effect of sex (F (1, 36) = 8.719, p = 0.0055), with males demonstrating greater TNF levels than females, this sex effect appeared to be driven by the exercised males, as there was a trend for the interaction between exercise and sex (F (1, 36) = 3.390, *p* = 0.0738), but no effect of exercise (F (1, 36) = 0.6252, *p* = 0.4343).

**Figure 4.**
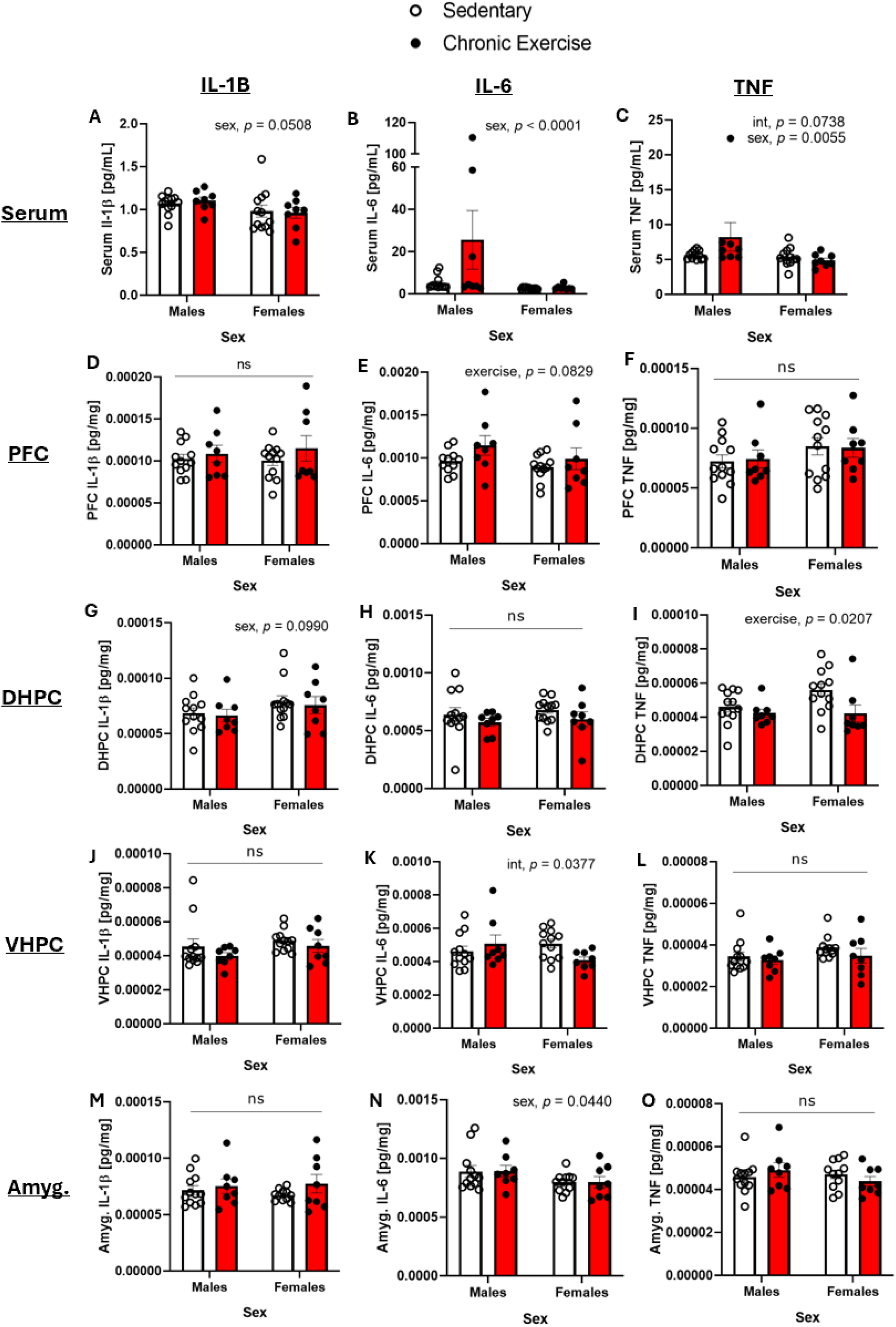
Effect of chronic exercise and sex on IL-1β, IL-6 and TNF. Serum levels of (A) IL-1β, (B) IL-6, and (C) TNF. Prefrontal cortical levels of (D) IL-1β, (E) IL-6) and (F) TNF. Dorsal hippocampal levels of (G) IL-1β, (H) IL-6, and (I) TNF. Ventral hippocampal levels of (J) IL-1β, (K) IL-6, and (L) TNF. Amygdala levels of (M) IL-1β, (N) IL-6, and (O) TNF. Concentration levels were adjusted to protein concentration and are expressed as pg. cytokine/mg. protein. Error bars indicate standard error. Note, * p < 0.05. n = 11-12 per sex in sedentary group and n = 7-8 per sex in exercised group.

#### Specific changes in brain cytokines after chronic exercise

In the prefrontal cortex (Figure 4D-F), there was no effects of exercise, sex, or interaction on levels of IL-1β (exercise (F (1, 36) = 1.382, p = 0.2475), sex (F (1, 36) = 0.06356, p = 0.8024), or interaction (F (1, 36) = 0.2252, *p* = 0.6379, Figure 4D) or TNF (exercise: (F (1, 36) = 0.002641, p = 0.9593), sex: (F (1, 36) = 2.426, p = 0.1281), interaction: (F (1, 36) = 0.05412, p = 0.8174). For IL-6, there was a trend with exercised animals having higher IL-6 levels compared to sedentary animals (*F* (1, 35) = 3.186; p = 0.0829), but there was no effect of sex (F (1, 35) = 2.054, p = 0.1607), nor an interaction (F (1, 35) = 0.2996, p = 0.5876); Figure 4E).

In the dorsal hippocampus (Figure 4G-I), there was an effect of chronic exercise on TNF levels, with exercised animals demonstrating lower concentrations of this cytokine (F (1, 36) = 5.859, p = 0.0207) in both sexes (no significant interaction (F (1, 36) = 1.976, p = 0.1684), or main effect of sex (F (1, 36) = 1.886, p = 0.1781; Figure 4I). There was no effect of exercise, sex, or interaction on dorsal hippocampal IL-1β (exercise: (F (1, 36) = 0.2015, p = 0.6562), sex: (F (1, 36) = 2.868, p = 0.0990), interaction: (F (1, 36) = 0.01006, p = 0.9207) and IL-6 levels (exercise: (F (1, 36) = 2.167, p = 0.1497), sex: (F (1, 36) = 0.4450, p = 0.5090), interaction: (F (1, 36) = 0.01622, p = 0.8994.

In the ventral hippocampus (Figure 4J-L), there was an interaction for IL-6 (F (1, 36) = 4.654, p = 0.0377), but follow up comparison tests revealed only a trend, with exercised females showing lower IL-6 levels than the sedentary females (p = 0.0793). No difference was observed between sedentary and exercised males (p = 0.5917). There was no effect of chronic exercise (F (1, 36) = 0.7206, p = 0.4016), or sex (F (1, 36) = 0.6447, p = 0.4273) on IL-6 levels. There was also no effect of chronic exercise, sex, or interaction on ventral hippocampal IL-1β (exercise: (F (1, 36) = 1.649, p = 0.2074), sex: (F (1, 36) = 1.917, p = 0.1747), interaction: (F (1, 36) = 0.1412, p = 0.7093), or TNF levels (exercise: (F (1, 36) = 1.429, p = 0.2397), sex: (F (1, 36) = 1.836, p = 0.1838): interaction: (F (1, 36) = 0.2441, p = 0.6242).

In the amygdala (Figure 4M-O), there was no effect of exercise, sex, or interaction for IL-1β (exercise: (*F* (1, 35) = 1.795, p = 0.1890), sex: (*F* (1, 35) = 0.04363, *p* = 0.8357), interaction: (*F* (1, 35) = 0.4131, *p* = 0.5246)) or TNF (exercise: (F (1, 35) = 0.0001239, p = 0.9912), sex: (F (1, 35) = 0.6452, p = 0.4273), interaction: (F (1, 35) = 1.757, p = 0.1936)). There was an effect of sex on IL-6 levels, with females exhibiting lower IL-6 than males (*F* (1, 35) = 4.367, *p* = 0.0440). There was no main effect of exercise (*F* (1, 35) = 0.0002977, *p* = 0.9863), and no interaction (*F* (1, 35) = 0.0007966, *p* = 0.9776).

#### Characterization of patterns in serum cytokines after chronic exercise

A principal component analysis of all cytokine concentrations in the serum of the chronic exercise cohort (Supplementary Figure 6A) revealed that three factors accounted for 67.19% of biomarker variation (Chronic Serum Factor 1, 34.01%; Chronic Serum Factor 2, 19.58%; Chronic Serum Factor 3, 13.59%). To test for exercise and sex-dependent differences in the mean values of these extracted components, three separate ANOVAs were performed on Chronic Serum Factor 1, 2 and 3. For Chronic Serum Factor 1, there was a significant effect of sex (F (1, 36) = 5.800, p = 0.0213), with males demonstrating higher measures of Chronic Serum Factor 1 (driven by high loadings of KC/GRO, IL-6 and IL-1β; Supplementary Figure 6B). Although this sex effect appeared to be driven by the exercised males, there was no effect of exercise (F (1, 36) = 1.920, p = 0.1743), nor an interaction (F (1, 36) = 1.412, p = 0.2425). For Chronic Serum Factor 2, there was no effect of exercise (F (1, 36) = 1.147, p = 0.2914), sex (F (1, 36) = 0.004951, p = 0.9443), or an interaction (F (1, 36) = 0.1638, p = 0.6880; Supplementary Figure 6C)). For Chronic Serum Factor 3, there was a significant effect of sex (F (1, 36) = 43.49, p = 0.0001), with females demonstrating higher measures of Chronic Serum Factor 3 than males (driven by high loadings of IL-5 and negative loadings of IL-6; Supplementary Figure 6D). However, there was no effect of exercise (F (1, 36) = 0.02229, p = 0.8821), nor an interaction (F (1, 36) = 0.4362, p = 0.5131).

#### Characterization of patterns in brain cytokines after chronic exercise

A principal component analysis of all cytokine concentrations within the prefrontal cortex of the chronic exercise cohort (Supplementary Figure 7A) revealed that three factors accounted for 71.80% of biomarker variation (Chronic PFC Factor 1, 43.84%, Chronic PFC Factor 2, 16.67%, Chronic PFC Factor 3, 11.30%). For Chronic PFC Factor 1, there was no effect of exercise (*F* (1, 36) = 1.600, *p* = 0.2141), or interaction (F (1, 36) = 0.1261, p = 0.7245), but there was a significant effect of sex (*F* (1, 36) = 4.270, *p* = 0.0460), with males demonstrating higher measures of Chronic PFC Factor 1 (driven by high loadings of IL-6, IFN-γ, IL-12p70, and IL-5; Supplementary Figure 7B). There was also a near significant effect of sex for Chronic PFC Factor 2, with a trend for females demonstrating higher measures of Chronic PFC Factor 2 (*F* (1, 36) = 3.549, *p* = 0.0677), but there was no effect of exercise (*F* (1, 36) = 0.4603, *p* = 0.5018), or interaction (*F* (1, 36) = 0.9596, *p* = 0.3338; Supplementary Figure 7C)). Finally, there was no effect of exercise (*F* (1, 36) = 0.007974, *p* = 0.9293), sex (F (1, 36) = 0.1487, p = 0.7020), or an interaction (*F* (1, 36) = 0.1006, *p* = 0.7529) for Chronic PFC Factor 3 (Supplementary Figure 7D).

In the dorsal hippocampus, a principal component analysis of cytokine concentrations (Supplementary Figure 8A) revealed that two factors accounted for 63.93% of biomarker variation (Chronic DHPC Factor 1,50.32%, Chronic DHPC Factor 2, 13.61%). For Chronic DHPC Factor 1 (Supplementary Figure 8B), there was no effect of exercise (F (1, 36) = 3.055, p = 0.089), sex (F (1, 36) = 1.303, p = 0.2612), or interaction (F (1, 36) = 3.013, p = 0.0912). For Chronic DHPC Factor 2 (Supplementary Figure 8C), there was an effect of sex (F (1, 36) = 11.37, p = 0.0018; driven by high loadings of IL-5, IL-2, and IL-1β), but no effect of exercise (F (1, 36) = 0.9150, p = 0.3452), or interaction (F (1, 36) = 0.4328, p = 0.5148; Supplementary Figure 8C).

Analysis of ventral hippocampal cytokine concentrations using principal component analysis (Supplementary Figure 9A) revealed two principal factors that together explained 70.87% of the overall variance (Chronic VHPC Factor 1, 59.73%, Chronic VHPC Factor 2, 11.14%). There was no effect of exercise (F (1, 36) = 1.705, p = 0.1999), sex (F (1, 36) = 0.02202, p = 0.8829), or interaction (F (1, 36) = 0.3139, 0.5788) for Chronic VHPC Factor 1 (Supplementary Figure 9B). For Chronic VHPC Factor 2, there was an effect of sex, with females having higher measures of Chronic VHPC Factor 2 than males (F (1, 36) = 4.263, p = 0.0462; driven by high loadings of IL-12p70 and IL-2). However, there was no effect of exercise (F (1, 36) = 0.2245, p = 0.6385), nor an interaction (F (1, 36) = 0.05092, p = 0.8227) for Chronic VHPC Factor 2 (Supplementary Figure 9C).

In the amygdala, a principal component analysis of all cytokine concentrations (Supplementary Figure 10A) revealed that three factors accounted for 72.76% of biomarker variation (Chronic Amygdala Factor 1, 49.95%, Chronic Amygdala Factor 2, 11.82%, Chronic Amygdala Factor 3, 10.98%). For Chronic Amygdala Factor 1, there was no effect of exercise (F (1, 35) = 0.1331, p = 0.7174), sex (F (1, 35) = 1.668, p = 0.2050) or an interaction (F (1, 35) = 0.09366, p = 0.7614; Supplementary Figure 10B). Likewise, there was no effect of exercise (F (1, 35) = 0.2664, p = 0.6090), sex (F (1, 35) = 0.07772, p = 0.7821), nor an interaction (F (1, 35) = 0.6512, p = 0.4251) for Chronic Amygdala Factor 2 (Supplementary Figure 10C), or Chronic amygdala Factor 3 (exercise, F (1, 35) = 1.002, p = 0.3236; sex, F (1, 35) = 0.9478, p = 0.3370; interaction, F (1, 35) = 0.4137, p = 0.5243; Supplementary Figure 10D).

## DISCUSSION

This study provides new insights into the distinct effects of acute and chronic exercise on anxiety-related behaviours and circulating and brain cytokine changes in healthy, young adult male and female mice. Key findings demonstrate that acute exercise increased anxiety-like behaviour in males, and altered circulating cytokine levels in both sexes, marked by higher serum IL-6 and lower TNF, and an altered cytokine profile in the amygdala distinguished by the higher levels of Acute Amygdala Factor 1 (driven by high loadings of IL-12p70, IL-10, IFN-γ, and TNF). Chronic exercise was found to reduce anxiety-like behaviour in both sexes and lower TNF content specifically in the dorsal hippocampus, with no differences observed in circulating cytokines. These findings underscore the potential time-dependent impact of exercise of moderate intensity and sufficient duration on immune markers and provide insight into the underlying mechanisms that promote cognitive health and well-being across sex with sustained exercise.

### Acute exercise increased anxiety-like behaviour in males

Based on previous work in humans (21) and mixed previous work in rodents (9–11), it was predicted that acute exercise would reduce anxiety-like behaviour in both sexes, with a potentially greater reduction in females. Contrary to expectations, exercise decreased average speed and increased time spent immobile in both sexes. Additionally, exercised males spent less time in the centre of the arena compared to sedentary males. Accordingly, in contrast to our hypothesis, these results suggest that acute exercise increased anxiety-like behaviour, particularly in males more than females. Our exercise paradigm was moderate in intensity and the mice recovered for two hours post-exercise, prior to conducting behavioural assays to minimize the effects of exhaustion. Our results are consistent with previous work conducted in adult male C57BL/6J mice, wherein acute treadmill running at moderate (12 m/min) and high (15-17 m/min) intensities and for durations of 30 minutes has been demonstrated to decrease exploratory behaviour and increase grooming behaviour immediately following exercise (10). Given that measures of anxiety are often related to locomotion, and that locomotion can be confounded by exhaustion in exercise contexts, it is notable that exercised animals demonstrated reduced locomotion and time in the centre even two hours after moderate treadmill running. Additionally, although changes in speed and immobility could reflect exercise-induced exhaustion rather than anxiety, the number of entries made into the centre of the arena was not affected by exercise, which further controverts exhaustion. Interestingly, a low-speed treadmill running appears to reduce anxiety- and depressive-like behaviour in male rats, measured 30-minutes post exercise, in the elevated plus maze and forced swim test, which was not observed using the same exercise paradigm at higher speeds (9). However, a difficulty of using acute treadmill exercise in rodent models pertains to the potentially stressful nature of the intervention. Although exercise generally activates the HPA axis, this activation may be more prevalent in acute forced exercise regimes, as recent work has shown that acute forced treadmill exercise elicits activation of brain areas associated with pain, fear, and stress (46). Compared to running wheel, treadmill exposure alone appears to more strongly activate the paraventricular nucleus of the hypothalamus, and increase corticosterone levels in mice (46). Consequently, acute forced exercise may be perceived as more stressful than voluntary paradigms, thereby posing a confounding factor on behavioural outcomes. Overall, our study contributes to the growing literature on the effects of acute treadmill exercise on anxiety-like behaviour in healthy male and female rodents. Future work should continue to develop a systematic approach to elucidating the behavioural effects of acute treadmill exercise while further minimizing confounding factors such as stress and exhaustion.

### Acute exercise regulates serum and amygdala cytokines

In this study, we used both targeted and exploratory approaches to elucidate changes in various cytokines and chemokines after exercise. In our targeted approach, we specifically analyzed stress and exercise-responsive cytokines: IL-1β, IL-6 and TNF. Two hours post-acute exercise, serum IL-6 levels were higher, while TNF levels were lower in both male and female mice compared to sedentary animals. IL-6 acts as a primary mediator of the exercise-induced response to exercise (47,48). It is rapidly increased in serum with acute exercise and has been shown to regulate the levels of other cytokines, including TNF and IL-1β, although this has been primarily demonstrated in human males (14). The elevated serum IL-6 observed in this study is consistent with previous work demonstrating post-exercise increases in IL-6 across multiple species, but mostly in males (14,23,49–51). However, limited work has shown this increase in IL-6 in circulation in healthy male mice (36,52), with previous work focusing on other tissues in mostly male rodents, such as muscle (53), adipose (52), and intestinal lymphocytes (37). Additionally, the literature on TNF responses to acute exercise are mixed: previous work has demonstrated an increase in serum TNF in male humans (54,55), while others have demonstrated a decrease (56) and no change in male mice (49). These discrepancies likely reflect differences in the type, duration, intensity of the exercise, as well as timing of blood sampling, and the animal being studied. Finally, although IL-1β did not change with acute exercise, this may be explained by previous work demonstrating increases primarily in strenuous exercise contexts (54,57,58) and by the limited work measuring this cytokine in rodent exercise paradigms (37). These results show that the transient exercise-induced changes in serum cytokines did not correspond with parallel changes in brain cytokines, as no changes in IL 1β, IL-6, or TNF were detected in any brain region analyzed. The only central change observed with acute exercise was an increase in Acute Amygdala Factor 1, characterized by high loadings of IL-12p70, IL-10, IFN-γ, and TNF. Previous work has shown that forced exercise paradigms, such as a treadmill running, can activate brain areas associated with pain and fear in male mice (46). Accordingly, the heightened levels of Acute Amygdala Factor 1 observed in the current study may reflect activation in the amygdala, potentially associated with heightened anxiety. Altogether, these results demonstrate changes in serum cytokines following an acute bout of exercise, consistent with previous human literature (14,56), but more nuanced changes in the profile of central cytokines at 2-hours post-exercise.

Finally, a key question is why both sexes exhibited similar exercise-induced changes in immune signaling despite sex differences in anxiety-related behaviour following exercise. It is possible that changes in amygdala cytokines do not similarly regulate behaviour in both sexes. Another possibility is that anxiety may be influenced by hormone levels. Indeed, the female estrous cycle is associated with changes in anxiety-like behaviour in mice (59,60). However, how hormone levels in both sexes, estrous cycle, and exercise interact to regulate anxiety is not clear. Future studies should investigate the role of hormones and estrous cycle on behavioural outcomes with respect to exercise.

### Chronic exercise decreased anxiety-like behaviour in both sexes

The chronic exercise regimen increased the number of entries made into the centre of the open field and reduced grooming behaviour in both sexes. Increased self-grooming can be an indicator of stress (61), and increased time spent in the centre of the open field is often associated with reduced anxiety (62). Chronic exercise also increased time spent immobile among females, which has also been reported previously (63). Although increased immobility may be an indicator of increased anxiety, the preference for immobility may have been an active coping mechanism facilitating centre exploration, given that exercised mice had more centre entries, and the centre is a more anxiety-provoking region that may yield immobility. Recent work has shown that in male Sprague Dawley rats, 30 minutes of daily low-intensity treadmill running for eight weeks reduced anxiety-like behaviour, even when a foot shock was applied to encourage running (64). When considered with the grooming and location data, our results indicate that chronic exercise reduced anxiety-like behaviour in both sexes, which is consistent with previous work investigating treadmill exercise, stress coping, and anxiety in rodents of either sex (63–65), as well as previous research using running wheel paradigms in female mice (66).

### Chronic exercise minimally altered serum cytokines and decreased TNF in the dorsal hippocampus

Based on previous research regarding the anti-inflammatory effects of chronic exercise in humans (23), it was predicted that chronic exercise would decrease both serum and brain levels of IL-1β, IL-6, and TNF. Contrary to expectations, chronic exercise did not significantly alter these cytokines in the serum. However, the cytokine response in the chronic regimen differed from that of the acute, as the exercise effects observed from the acute bout were not sustained in the chronic regimen. These results indicate that despite a shift in cytokine responses from acute to chronic exercise, the latter may not significantly alter serum levels of IL-1β, IL-6, and TNF in a healthy model, despite previous work showing a shift to anti-inflammatory signaling in aged and disease models with routine exercise (26,27,67–69).

Despite negligible changes in serum cytokines, chronic exercise decreased TNF within the dorsal hippocampus in both sexes, and trended in decreasing chronic Dorsal Hippocampal Factor 1, which also loaded TNF among other cytokines (IL-12p70, IL-10, IFN-γ, KC/GRO, and IL-6). This result is in line with previous work in male mice that demonstrated decreased levels of hippocampal TNF following 16 weeks of voluntary wheel running (37). The decrease in TNF in the dorsal hippocampus may also further relate to the reduced anxiety outcomes observed among the chronic exercised group, as previous research indicates that limbic-derived TNF signaling is important for anxiety-like behaviour in mice (70). Overall, the chronic exercise-induced decreases observed in dorsal hippocampal TNF are consistent with previous work and in agreement with the observed decreases in anxiety-like behaviour.

### Sex differences in cytokine levels are present in circulation and brain

In addition to exercise effects, this study identified sex differences with respect to cytokine expression in the blood and brain. The inclusion of both sexes is an overall strength of this study, as females have been grossly understudied in both the field of neuroscience and the field of exercise physiology (71,72). Males had significantly higher IL-6 expression in the serum, prefrontal cortex, and dorsal hippocampus of compared to females, regardless of whether animals exercised or not. Although there has been limited work in healthy models demonstrating such differences, research in various states of disease and injury have demonstrated heightened IL-6 levels in the serum of male humans (73–76). Contrary to our findings, previous work found that adult male and female C57B/6N mice do not significantly differ in hippocampal IL-6 mRNA expression (77). These contradictory findings could be related to how the hippocampus was analyzed. The dorsal and ventral hippocampus have distinct functions, with the dorsal region being dominantly involved in spatial learning and memory, whereas the ventral region is more closely associated with stress reactivity and anxiety regulation (45,78). To explore the potential mechanisms of these bifunctional phenomena, we separated the dorsal and ventral hippocampus and found no sex-related IL-6 differences in the ventral region, as well as minimal individual cytokine-level sex differences in this region overall. Additionally, although we found several interaction effects of exercise and sex on various cytokines, these interactions were from sex-differences in the control groups, which were not present in the exercise groups. However, we did find sex-related differences in dorsal and ventral hippocampus factors, which were comprised of different cytokine loadings, thereby indicating systemic differences in subregional hippocampal cytokine expression. Although there is minimal research regarding sex differences in cytokine changes in the dorsal and ventral hippocampus following exercise, previous work has demonstrated that male and female mice have different inflammatory hippocampal responses to immune challenge (79) and stress (80). Contrary to stress and immune-challenge research, our work demonstrated that exercise-induced cytokine changes also occurred across sex. Accordingly, it is possible that the sexes respond similarly to acute and chronic aerobic exercise, despite baseline cytokine differences. Given the limited sex-specific exercise research, further work is required. Every sex difference that is elucidated, along with every sex similarity, represents a gainful impact in our understanding of sex-based exercise prescription.

### Conclusion and significance

To conclude, we found differential effects of acute versus chronic exercise on anxiety-like behaviour and inflammatory signaling in the circulation and brain of healthy young adult male and female mice. This study contributes to our understanding of how exercise affects behaviour and underlying biological mechanisms in both sexes. In addition, our work may help inform future work pertaining to sex-based exercise prescription in the context of preventative medicine.

## ACKNOWLEDGEMENTS

We thank the animal care staff at the Comparative Bioscience Facility at Brock University for technical assistance. A special thanks to Talia Buffone who assisted with behavioural scoring. This research was supported by the Natural Sciences and Engineering Council of Canada to RM (RGPIN-2017-03904) and PDG (RGPIN-2022-04824), and the Canada Research Chair Program to PDG (2020-00099). MM was funded by an NSERC Canada Graduate Scholarship-Masters (CGS-M) and Branch Out Neurological Foundation.

