## Supplemental Figures for "Can you run from your worries? The effects of exercise on anxiety-like behaviour and immune signaling in female and male mice"

A

| Cytokine | Acute Serum Factor 1 | Acute Serum Factor 2 | Acute Serum Factor 3 |
|---|---|---|---|
| KC/GRO | 0.892 | | 0.148 |
| IL-6 | 0.869 | | -0.286 |
| IL-10 | 0.657 | 0.126 | 0.183 |
| IL-1β | 0.582 | 0.142 | 0.416 |
| TNF | 0.301 | 0.368 | -0.227 |
| IL2 | 0.153 | 0.843 | 0.548 |
| IFN- γ | | 0.586 | 0.267 |
| IL-5 | | 0.518 | 0.788 |

| Legend |
|---|
| 0.9+ |
| 0.8-0.899 |
| 0.6-0.799 |
| 0.4-0.599 |
| 0.3-0.399 |
| (-0.299) - 0.299 |
| (-0.399) - (-0.3) |
| (-0.599) - (-0.4) |
| (-0.799) - (-0.6) |
| (-0.899) - (-0.8) |
| (-0.9)- |

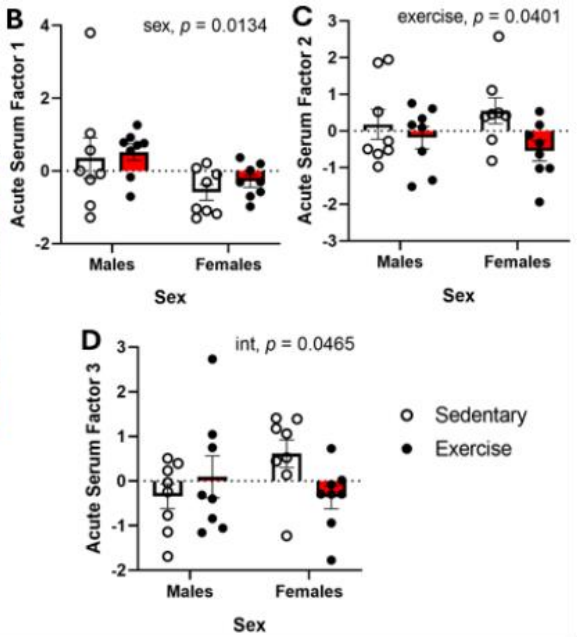


**Supplementary Figure 1. Principal component analysis of serum cytokines of the acute exercise cohort.** (A) Rotated component matrix loadings. Components were subject to varimax rotation with Kaiser normalization and converged in 5 iterations. Only loadings >0.1 are reported, and only loadings >0.3 may be considered meaningful. ANOVAs were performed on all principal components. (B) Females had lower measures for Acute Serum Factor 1 ($p < 0.05$). (C) Exercised mice had lower measures of Acute Serum Factor 2 ($p < 0.05$). (D) Multiple comparison tests showed no significant differences with respect to the significant interaction effect for Acute Serum Factor 3. Statistical details are described in text.

A

| Cytokine | Acute PFC Factor 1 | Acute PFC Factor 2 |
|---|---|---|
| IL-2 | 0.916 | |
| IFN- γ | 0.911 | 0.160 |
| TNF | 0.874 | |
| IL-5 | 0.862 | 0.198 |
| IL-6 | 0.861 | |
| IL-1β | 0.817 | |
| IL-12p70 | 0.807 | 0.336 |
| IL-10 | 0.742 | 0.517 |
| KC/GRO | 0.712 | |
| IL-4 | | 0.964 |

| Legend |
|---|
| 0.9+ |
| 0.8-0.899 |
| 0.6-0.799 |
| 0.4-0.599 |
| 0.3-0.399 |
| (-0.299) - 0.299 |
| (-0.399) - (-0.3) |
| (-0.599) - (-0.4) |
| (-0.799) - (-0.6) |
| (-0.899) - (-0.8) |
| (-0.9)– |

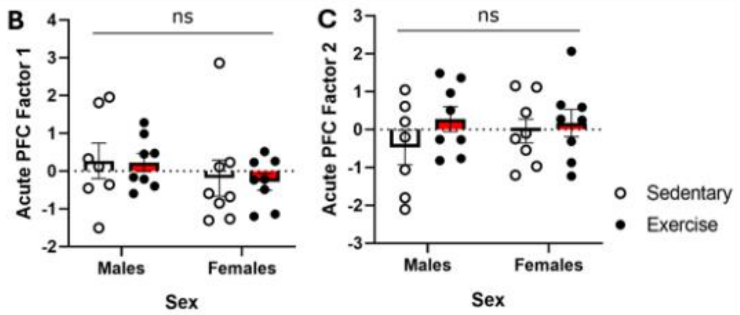


**Supplementary Figure 2. Principal component analysis of prefrontal cortex cytokines of the acute exercise cohort.** (A) Rotated component matrix loadings. Components were subject to varimax rotation with Kaiser normalization and converged in 3 iterations. Only loadings >0.1 are reported, and only loadings >0.3 may be considered meaningful. ANOVAs were performed on both principal components and revealed no significant exercise or sex effects were for (B) Acute PFC Factor 1 or (C) Acute PFC Factor 2. Statistical details are described in text.

A

| Cytokine | Acute DHPC Factor 1 | Acute DHPC Factor 2 | Acute DHPC Factor 3 |
|---|---|---|---|
| IL-5 | 0.880 | 0.222 | |
| IL-10 | 0.782 | 0.252 | 0.166 |
| Il-2 | 0.729 | -0.116 | 0.370 |
| TNF | 0.687 | 0.249 | -0.315 |
| IFN- γ | 0.620 | 0.317 | 0.332 |
| IL-1β | 0.589 | | |
| IL-6 | 0.525 | 0.497 | -0.472 |
| KC/GRO | | 0.945 | -0.122 |
| IL-12p70 | 0.542 | 0.632 | |
| IL-4 | 0.181 | 0.167 | 0.883 |

| Legend |
|---|
| 0.9+ |
| 0.8-0.899 |
| 0.6-0.799 |
| 0.4-0.599 |
| 0.3-0.399 |
| (-0.299) - 0.299 |
| (-0.399) - (-0.3) |
| (-0.599) - (-0.4) |
| (-0.799) - (-0.6) |
| (-0.899) - (-0.8) |
| (-0.9)– |

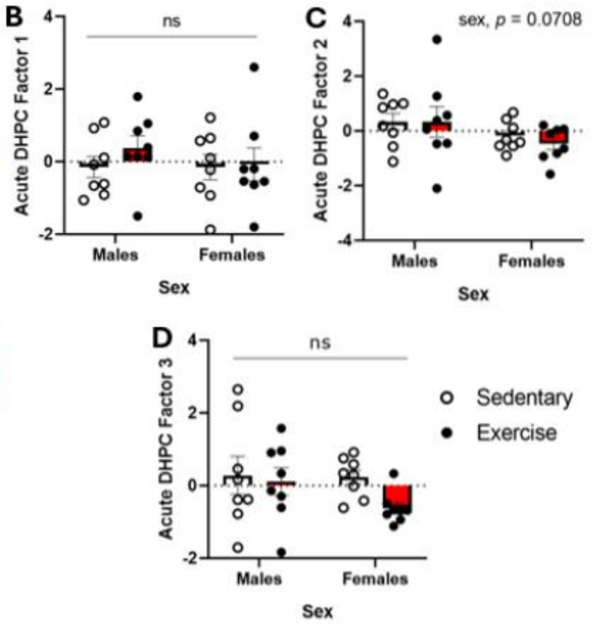


**Supplementary Figure 3. Principal component analysis of dorsal hippocampus cytokines of the acute exercise cohort.** (A) Rotated component matrix loadings. Components were subject to varimax rotation with Kaiser normalization and converged in 8 iterations. Only loadings >0.1 are reported, and only loadings >0.3 may be considered meaningful. ANOVAs were performed on all principal components and revealed no significant exercise or sex effects for (B) Acute DHPC Factor 1, (C) Acute DHPC Factor 2, or (D) Acute DHPC Factor 3. Statistical details are described in text.

**A**

| Cytokine | Acute VHPC Factor 1 | Acute VHPC Factor 2 | Acute VHPC Factor 3 | Acute VHPC Factor 4 |
|---|---|---|---|---|
| IL-12p70 | 0.885 | | -0.193 | |
| IL-5 | 0.844 | 0.198 | 0.158 | -0.167 |
| Il-2 | 0.696 | 0.390 | 0.188 | -0.261 |
| IL6 | 0.695 | -0.269 | 0.392 | 0.412 |
| IFN- γ | 0.617 | 0.259 | 0.601 | |
| IL-4 | | 0.916 | | |
| IL-10 | 0.294 | 0.661 | | 0.425 |
| IL-1β | | | 0.924 | |
| KC/GRO | | | 0.200 | 0.813 |
| TNF | 0.132 | -0.230 | 0.254 | -0.577 |

| Legend |
|---|
| 0.9+ |
| 0.8-0.899 |
| 0.6-0.799 |
| 0.4-0.599 |
| 0.3-0.399 |
| (-0.299) - 0.299 |
| (-0.399) - (-0.3) |
| (-0.599) - (-0.4) |
| (-0.799) - (-0.6) |
| (-0.899) - (-0.8) |
| (-0.9)– |

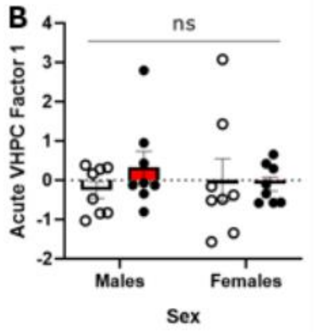


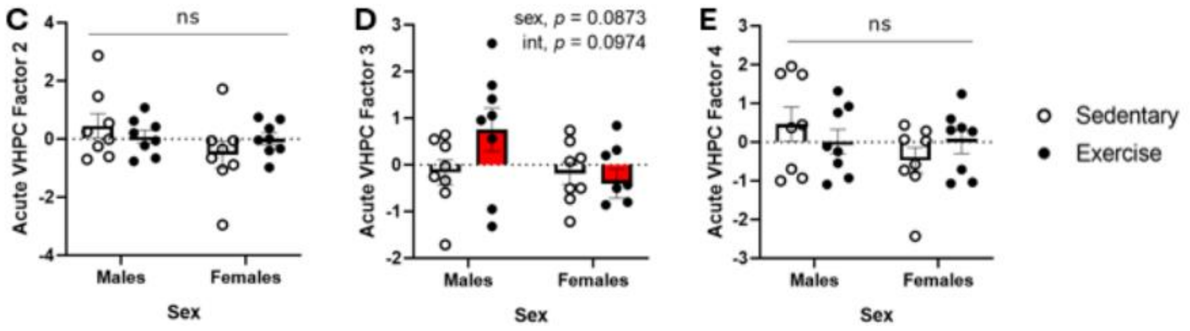


**Supplementary Figure 4. Principal component analysis of ventral hippocampus cytokines of the acute exercise cohort.** (A) Rotated component matrix loadings. Components were subject to varimax rotation with Kaiser normalization and converged in 7 iterations. Only loadings >0.1 are reported, and only loadings >0.3 may be considered meaningful. ANOVAs were performed on all principal components and revealed no significant exercise or sex effects for (B) Acute VHPC Factor 1, (C) Acute VHPC Factor 2, (D) Acute VHPC Factor 3, or (E) Acute VHPC Factor 4. Statistical details are described in text.

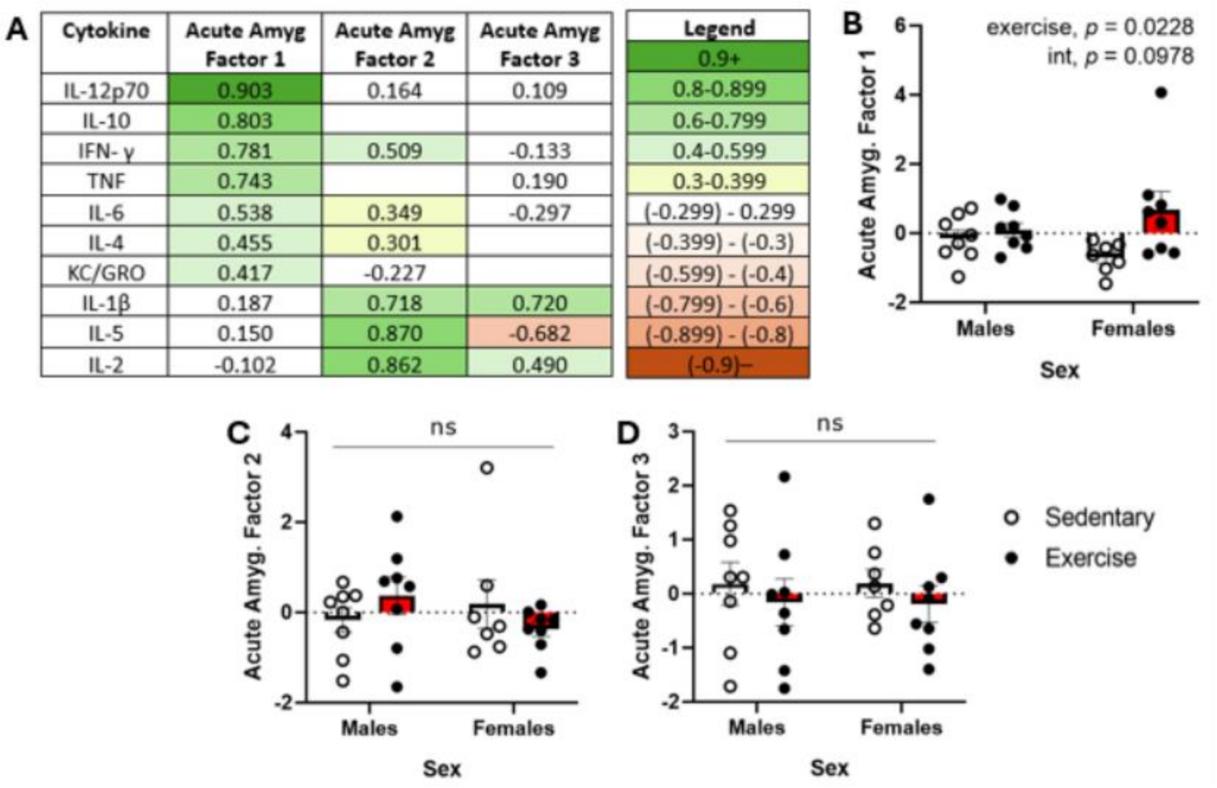

| Cytokine | Acute Amyg Factor 1 | Acute Amyg Factor 2 | Acute Amyg Factor 3 |
|---|---|---|---|
| IL-12p70 | 0.903 | 0.164 | 0.109 |
| IL-10 | 0.803 | | |
| IFN- γ | 0.781 | 0.509 | -0.133 |
| TNF | 0.743 | | 0.190 |
| IL-6 | 0.538 | 0.349 | -0.297 |
| IL-4 | 0.455 | 0.301 | |
| KC/GRO | 0.417 | -0.227 | |
| IL-1β | 0.187 | 0.718 | 0.720 |
| IL-5 | 0.150 | 0.870 | -0.682 |
| IL-2 | -0.102 | 0.862 | 0.490 |

| Legend |
|---|
| 0.9+ |
| 0.8-0.899 |
| 0.6-0.799 |
| 0.4-0.599 |
| 0.3-0.399 |
| (-0.299) - 0.299 |
| (-0.399) - (-0.3) |
| (-0.599) - (-0.4) |
| (-0.799) - (-0.6) |
| (-0.899) - (-0.8) |
| (-0.9)– |



**Supplementary Figure 5. Principal component analysis of amygdala cytokines of the acute exercise cohort.** (A) Rotated component matrix loadings. Components were subject to varimax rotation with Kaiser normalization and converged in 6 iterations. Only loadings >0.1 are reported, and only loadings >0.3 may be considered meaningful. ANOVAs were performed on all principal components and revealed a significant exercise effect for (B) Acute Amygdala Factor 1, but no exercise or sex effects for (C) Acute Amygdala Factor 2, or (D) Acute Amygdala Factor 3. Statistical details are described in text.

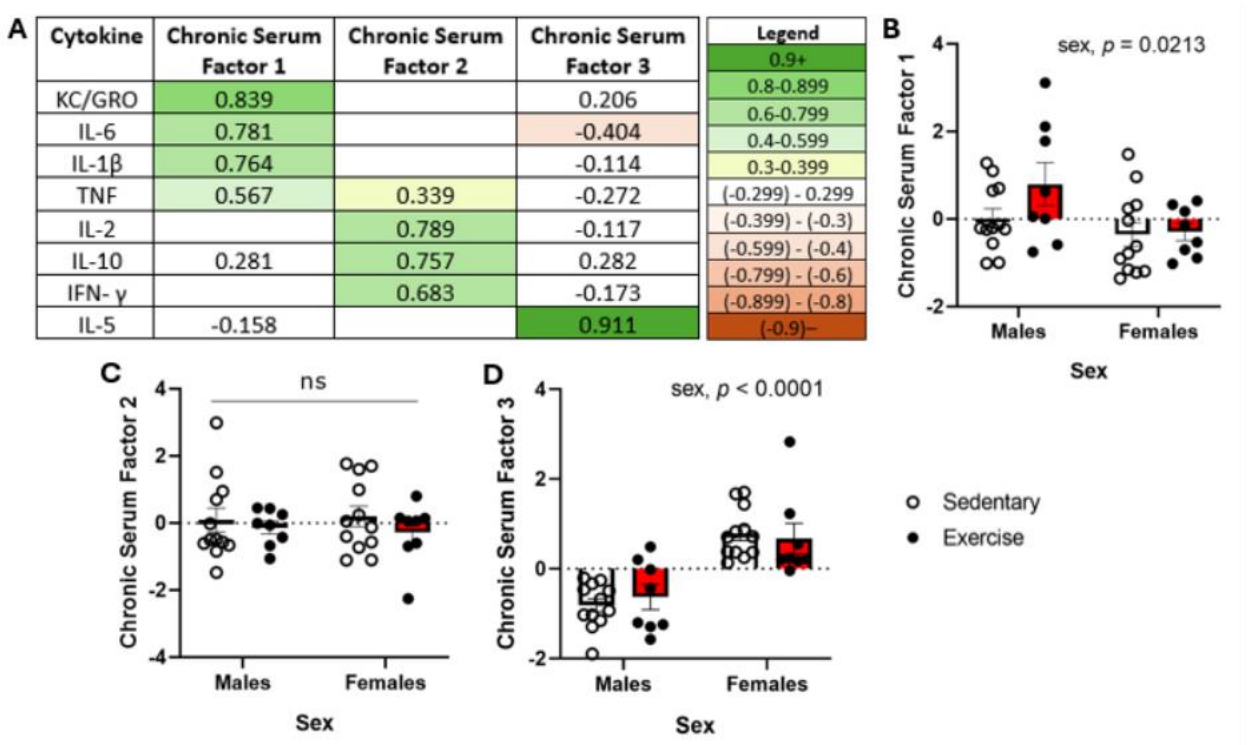

| Cytokine | Chronic Serum Factor 1 | Chronic Serum Factor 2 | Chronic Serum Factor 3 |
|---|---|---|---|
| KC/GRO | 0.839 | | 0.206 |
| IL-6 | 0.781 | | -0.404 |
| IL-1β | 0.764 | | -0.114 |
| TNF | 0.567 | 0.339 | -0.272 |
| IL-2 | | 0.789 | -0.117 |
| IL-10 | 0.281 | 0.757 | 0.282 |
| IFN- γ | | 0.683 | -0.173 |
| IL-5 | -0.158 | | 0.911 |



**Supplementary Figure 6. Principal component analysis of serum cytokines of the chronic exercise cohort.** (A) Rotated component matrix loadings. Components were subject to varimax rotation with Kaiser normalization and converged in 4 iterations. Only loadings >0.1 are reported, and only loadings >0.3 may be considered meaningful. ANOVAs were performed on all principal components and revealed a significant sex effect for (B) Chronic Serum Factor 1, no effects for (C) Chronic Serum Factor 2, and another sex effect for (D) Chronic Serum Factor 3. Statistical details are described in text.

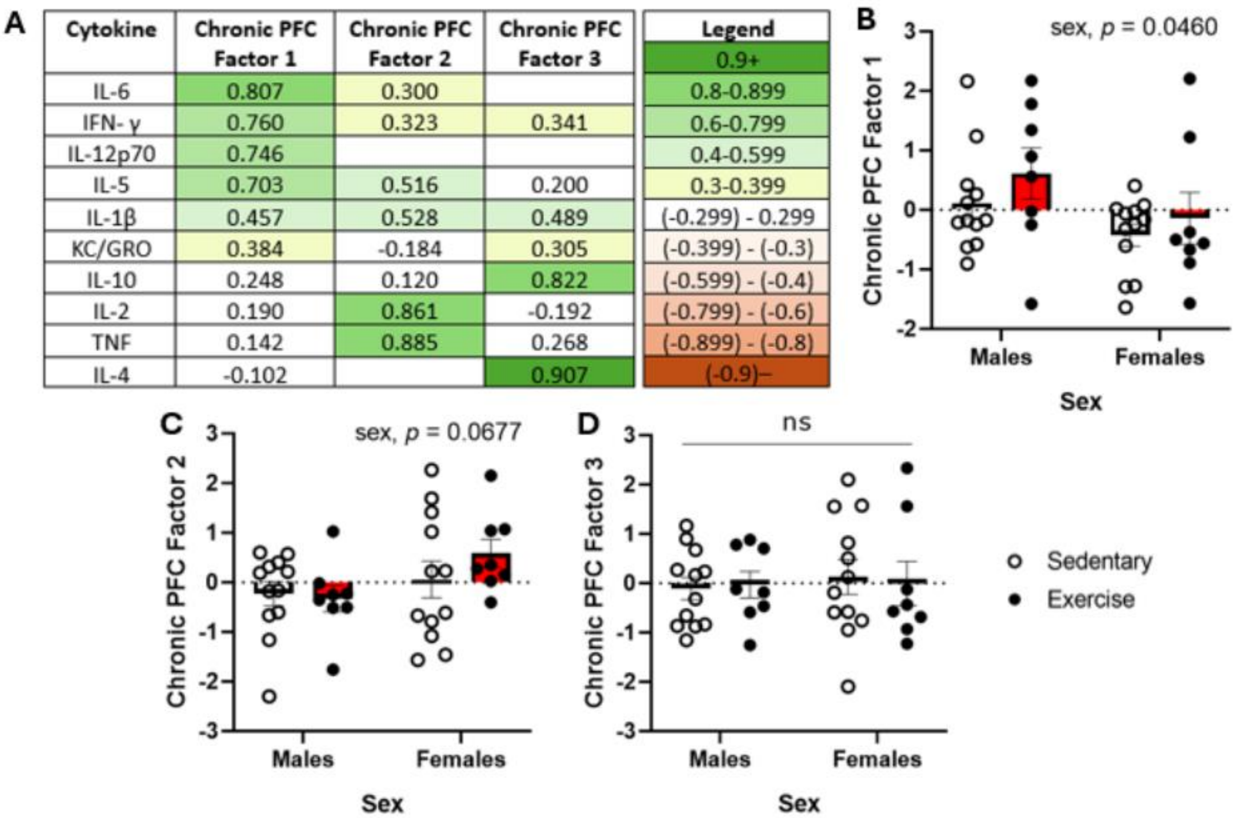

| Cytokine | Chronic PFC Factor 1 | Chronic PFC Factor 2 | Chronic PFC Factor 3 |
|---|---|---|---|
| IL-6 | 0.807 | 0.300 | |
| IFN- γ | 0.760 | 0.323 | 0.341 |
| IL-12p70 | 0.746 | | |
| IL-5 | 0.703 | 0.516 | 0.200 |
| IL-1β | 0.457 | 0.528 | 0.489 |
| KC/GRO | 0.384 | -0.184 | 0.305 |
| IL-10 | 0.248 | 0.120 | 0.822 |
| IL-2 | 0.190 | 0.861 | -0.192 |
| TNF | 0.142 | 0.885 | 0.268 |
| IL-4 | -0.102 | | 0.907 |

| Legend |
|---|
| 0.9+ |
| 0.8-0.899 |
| 0.6-0.799 |
| 0.4-0.599 |
| 0.3-0.399 |
| (-0.299) - 0.299 |
| (-0.399) - (-0.3) |
| (-0.599) - (-0.4) |
| (-0.799) - (-0.6) |
| (-0.899) - (-0.8) |
| (-0.9)– |



**Supplementary Figure 7. Principal component analysis of prefrontal cortex cytokines of the chronic exercise cohort.** (A) Rotated component matrix loadings. Components were subject to varimax rotation with Kaiser normalization and converged in 5 iterations. Only loadings >0.1 are reported, and only loadings >0.3 may be considered meaningful. ANOVAs were performed on all principal components and revealed a significant sex effect for (B) Chronic PFC Factor 1, and no significant effects for (C) Chronic PFC Factor 2, and (D) Chronic PFC Factor 3. Statistical details are described in text.

A

| Cytokine | Chronic DHPC Factor 1 | Chronic DHPC Factor 2 |
|---|---|---|
| IL-12p70 | 0.846 | 0.296 |
| IL-10 | 0.821 | 0.272 |
| TNF | 0.782 | 0.314 |
| IFN- γ | 0.751 | 0.223 |
| KC/GRO | 0.700 | |
| IL-6 | 0.475 | 0.563 |
| IL-1β | 0.235 | 0.650 |
| IL-2 | 0.153 | 0.759 |
| IL-5 | 0.141 | 0.865 |

| Legend |
|---|
| 0.9+ |
| 0.8-0.899 |
| 0.6-0.799 |
| 0.4-0.599 |
| 0.3-0.399 |
| (-0.299) - 0.299 |
| (-0.399) - (-0.3) |
| (-0.599) - (-0.4) |
| (-0.799) - (-0.6) |
| (-0.899) - (-0.8) |
| (-0.9)– |

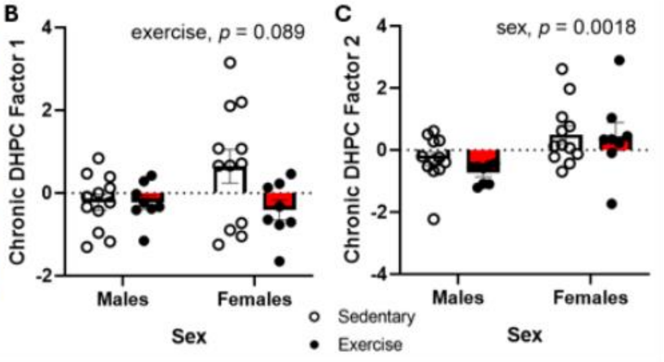


**Supplementary Figure 8. Principal component analysis of dorsal hippocampus cytokines of the chronic exercise cohort.** (A) Rotated component matrix loadings. Components were subject to varimax rotation with Kaiser normalization and converged in 3 iterations. Only loadings >0.1 are reported, and only loadings >0.3 may be considered meaningful. ANOVAs were performed on all principal components and revealed no significant exercise or sex effects for (B) Chronic DHPC Factor 1, and a significant sex effect for (C) Chronic DHPC Factor 2. Statistical details are described in text. Analyses excluded IL-4.

A

| Cytokine | Chronic VHPC Factor 1 | Chronic VHPC Factor 2 |
|---|---|---|
| IL-10 | 0.847 | 0.310 |
| IL-1β | 0.770 | 0.256 |
| KC/GRO | 0.747 | 0.108 |
| TNF | 0.736 | 0.524 |
| IL-5 | 0.725 | 0.481 |
| IL-6 | 0.713 | |
| IFN- γ | 0.603 | 0.630 |
| IL-2 | 0.411 | 0.734 |
| IL-12p70 | | 0.882 |

| Legend |
|---|
| 0.9+ |
| 0.8-0.899 |
| 0.6-0.799 |
| 0.4-0.599 |
| 0.3-0.399 |
| (-0.299) - 0.299 |
| (-0.399) - (-0.3) |
| (-0.599) - (-0.4) |
| (-0.799) - (-0.6) |
| (-0.899) - (-0.8) |
| (-0.9)– |

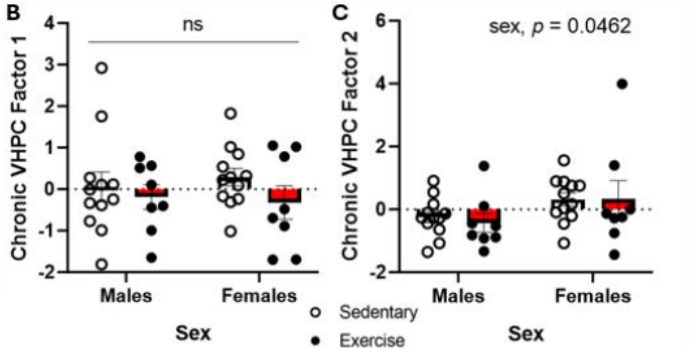


**Supplementary Figure 9. Principal component analysis of ventral hippocampus cytokines of the chronic exercise cohort.** (A) Rotated component matrix loadings. Components were subject to varimax rotation with Kaiser normalization and converged in 3 iterations. Only loadings >0.1 are reported, and only loadings >0.3 may be considered meaningful. ANOVAs were performed on both principal components and revealed no significant exercise or sex effects for (B) Chronic VHPC Factor 1, and a significant sex effect for (C) Chronic VHPC Factor 2. Statistical details are described in text. Analyses excluded IL-4.

A

| Cytokine | Chronic Amyg Factor 1 | Chronic Amyg Factor 2 | Chronic Amyg Factor 3 |
|---|---|---|---|
| IL-6 | 0.862 | 0.355 | 0.104 |
| IL-10 | 0.804 | 0.315 | 0.234 |
| IL-12p70 | 0.791 | 0.277 | |
| IL-5 | 0.767 | | 0.223 |
| IL-2 | 0.729 | 0.130 | -0.352 |
| TNF | 0.697 | | 0.345 |
| IFN- γ | 0.552 | 0.706 | |
| IL-1β | 0.348 | 0.547 | 0.273 |
| KC/GRO | 0.119 | | 0.908 |
| IL-4 | | -0.906 | |

| Legend |
|---|
| 0.9+ |
| 0.8-0.899 |
| 0.6-0.799 |
| 0.4-0.599 |
| 0.3-0.399 |
| (-0.299) - 0.299 |
| (-0.399) - (-0.3) |
| (-0.599) - (-0.4) |
| (-0.799) - (-0.6) |
| (-0.899) - (-0.8) |
| (-0.9)– |

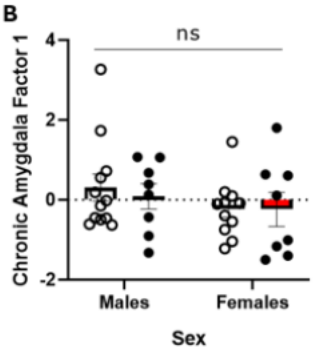


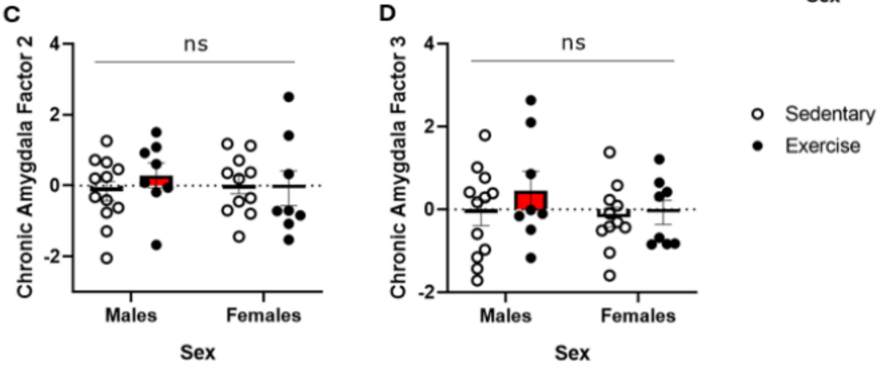


**Supplementary Figure 10. Principal component analysis of amygdala cytokines of the chronic exercise cohort.** (A) Rotated component matrix loadings. Components were subject to varimax rotation with Kaiser normalization and converged in 5 iterations. Only loadings >0.1 are reported, and only loadings >0.3 may be considered meaningful. ANOVAs were performed on all principal components and no significant effects for (B) Chronic Amygdala Factor 1, (C) Chronic Amygdala Factor 2, and (D) Chronic Amygdala Factor 3. Statistical details are described in text.
